# Cyb5r3-based mechanism and reversal of secondary failure to sulfonylurea

**DOI:** 10.1101/2022.03.23.485508

**Authors:** Hitoshi Watanabe, Wen Du, Jinsook Son, Lina Sui, Shun-ichiro Asahara, Irwin J Kurland, Taiyi Kuo, Takumi Kitamoto, Yasutaka Miyachi, Rafael de Cabo, Domenico Accili

## Abstract

Sulfonylureas (SU) are effective and affordable anti-diabetic drugs. But chronic use leads to secondary failure, limiting their utilization. The mechanism of secondary failure is unknown. Here we identify Cyb5r3 downregulation as a mechanism of SU failure and successfully reverse it. Chronic exposure to SU impairs Cyb5r3 levels and reduces islet glucose utilization with a metabolomics signature characterized by low acetyl-CoA and amino acid levels. Cyb5r3 engages in a glucose-dependent interaction that stabilizes glucokinase (Gck) to maintain glucose utilization. Accordingly, activating Gck mutations in patients with hyperinsulinemia reduce Cyb5r3 binding, whereas inactivating MODY mutations increase it, providing evidence for a role of Cyb5r3 in determining flux through Gck. The Cyb5r3 activator tetrahydroindenoindole (THII) rescues secondary failure to SU in an animal model of chronic SU treatment and restores insulin secretion from *ex vivo* islets. We conclude that Cyb5r3 loss-of-function is a key factor in the secondary failure to SU and a potential target for its prevention, which may lead to a rehabilitation of SU use in diabetes.

## Introduction

Impaired β-cell insulin secretion is a major etiological factor in type 2 diabetes. Sulfonylureas (SU) induce insulin secretion and, together with metformin, are commonly prescribed drugs for type 2 diabetes treatment worldwide ^1^. SU have been used for the management of type 2 diabetes since the 1950s and are still widely used not only in low- and middle-income countries, but also in developed countries because of their efficacy and low cost ^2, 3^. On the other hand, their use is often curtailed by secondary failure, risk of hypoglycemia, weight gain, and concerns about cardiovascular safety ^3^. Chronic treatment with SU is associated with secondary failure at a rate of 5-10% per year ^4–6^. In the UKPD study, after 6 years of SU treatment 20% of patients required combination therapy with metformin and 10% required insulin therapy^7^. The mechanism of secondary failure to SU is unknown ^8^.

SU promote insulin secretion by binding to the SU receptor 1 (SUR1), the regulatory subunit of the ATP-sensitive potassium (K_ATP_) channel, causing it to close, with ensuing depolarization, opening of voltage-dependent Ca2^+^ channels (VDSS), and influx of extracellular Ca2^+^ that triggers exocytosis of readily-releasable insulin granules ^9, 10^. As Cryo-EM studies have identified energy state-dependent glibenclamide (GLB), a prototypical third generation long-acting SU, binding sites in SUR1 that are inaccessible in low energy states ^11, 12^, Intracellular energy state likely to be closely related to SU responsiveness.

We have proposed that altered mitochondrial substrate utilization, or “metabolic inflexibility”, is an initial step in β-cell dedifferentiation, leading to β-cell failure ^13^. Transcription factors FoxO1 and FoxO3a are central to this model. An early manifestation of this process is the failure of GLB-induced insulin secretion along with glucose-induced insulin secretion ^13^. β-cell-specific FoxO1, FoxO3a, and FoxO4 knockout (the latter being hardly expressed in this cell type) promotes lipid over carbohydrate utilization as an energy source for oxidative phosphorylation, impairs ATP production, and lowers insulin release. Among the transcriptional targets of FoxO1, β-cell loss-of-function of the oxidoreductase cytochrome b5 reductase 3 (Cyb5r3), a component of the mitochondrial electron transfer system, affects glucose-induced insulin secretion, mitochondrial morphology, and ATP production ^14, 15^. The partial phenocopying of the FoxO1 knockout by the Cyb5r3 knockout raises the question of whether Cyb5r3 is the mediator of the early metabolic abnormalities leading to β-cell failure, including the failure of the SU response.

Here, we develop an *in vivo* model of SU failure and show that chronic GLB administration reduces protein levels of FoxO1, Cyb5r3 and glucokinase (Gck) as it impairs glucose-induced insulin secretion in human and mouse islets. We identify Cyb5r3 as a key regulator of SU responsiveness through direct, glucose-dependent binding to Gck that regulates the latter’s levels. Cyb5r3 inhibition results in constitutive failure of the GLB response that can be reversed either by a chemical Gck activator (GCKA) or, in the SU secondary failure model, by a Cyb5r3 activator.

## Results

### Molecular signature of SU secondary failure

As we have shown previously that FoxO1 ablation impairs the SU response, we evaluated FoxO1 localization in β-cells following prolonged exposure to GLB using islets from mice homozygous for a FoxO1-GFP knock-in allele (FoxO1-Venus) to facilitate its detection ^16^. We stimulated islets with GLB and monitored GFP localization for 96 h (Fig. 1a). In control islets, GFP localized to the cytoplasm from 24 to 96 h after stimulation. In contrast, in the GLB-treated group, GFP first underwent translocation to the nucleus (24 h), then to the cytoplasm (48 h), and finally disappeared (96 h). Western blotting confirmed that FoxO1 levels decreased after 96 h of GLB treatment (Fig 1b). Concurrently, islet GLB-induced insulin secretion decreased following 96 h treatment compared to vehicle-treated controls (Fig. 1c). To investigate the functional correlates of chronic GLB treatment on FoxO1 levels and insulin secretion in vivo, we developed an SU secondary failure model by implanting mice with a GLB-filled osmotic pump for up to 4 weeks and measured insulin and glucose levels. There was no difference in weight during the study period (Fig. S1a). Plasma insulin concentrations increased during the first week of GLB administration, but returned to basal during the second week, and tended to decrease thereafter (Fig. S1b,c). Glucose levels decreased after one week of GLB treatment but rebounded thereafter (Fig. S1d,e). These results suggest that the insulin secretory effect of GLB decreases after 2 weeks of chronic treatment, reminiscent of human data ^4^. Therefore, we analyzed GLB-induced insulin secretion after 3 weeks, and FoxO1 levels in isolated islets after 4 weeks of GLB administration. FoxO1 levels decreased (Fig. 1d), as did GLB-induced insulin secretion in mice treated with continuous GLB compared with controls, and the hypoglycemic effect of GLB was offset (Fig. 1e,f).

**Fig. 1.**
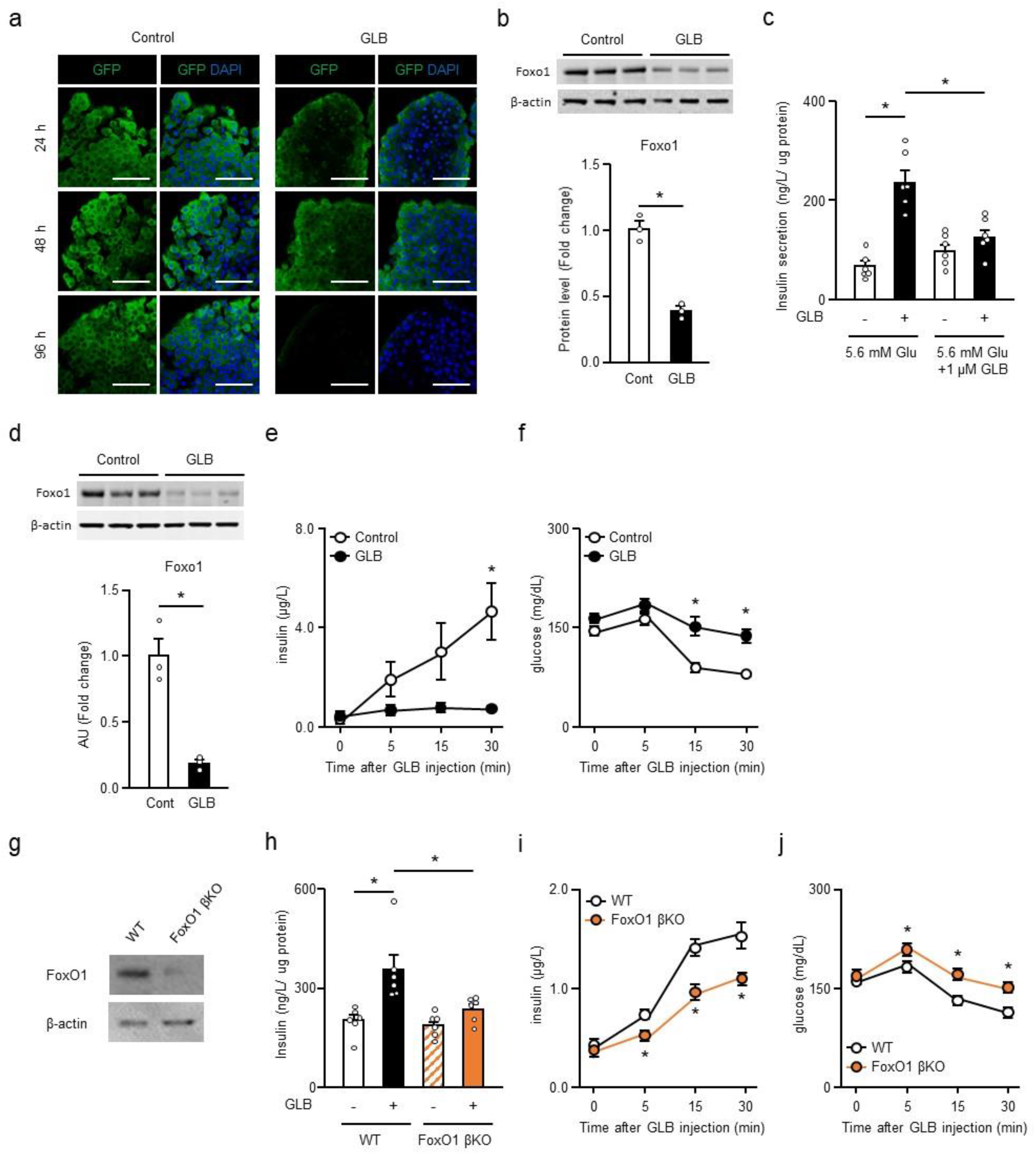
Chronic GLB decreases Foxo1 levels and GLB-induced insulin secretion. **a**, GFP immunostaining of islets from FoxO1-Venus mice following GLB treatment for 24 to 96 h. Scale bar: 50 μm. **b**, FoxO1 levels in islets with or without GLB treatment for 96 h (n=3). **c**, GLB-induced insulin secretion in islets with or without GLB treatment for 96 h (n=6). **d**, FoxO1 levels in islets from mice with or without GLB treatment for 4 weeks (n=3). **e,f**, Insulin (**e**) and glucose levels (**f**) after GLB administration in mice with or without GLB treatment for 3 weeks (n=6-7). **g**, FoxO1 levels in wild type (WT) and FoxO1 β-cell-specific knockout (FoxO1 βKO) islets. **h**, GLB-induced insulin secretion in islets from WT and FoxO1 βKO (n=6). **i,j**, Insulin (**i**) and glucose (**j**) levels after GLB administration in WT and FoxO1 βKO (n=5). \**P* < 0.05 by unpaired Student’s *t*-test and two-way ANOVA followed by Tukey’s test. Data represent means ± SEM.

To investigate whether the decrease of FoxO1 induced by chronic GLB administration is mechanistically involved in the failure of chronic GLB treatment, we analyzed the effects of GLB on insulin secretion using FoxO1 β-cell-specific knockout mice (FoxO1 βKO). We previously showed that GLB- induced insulin secretion is impaired in FoxO1/3a/4 triple knockout mice ^13^. GLB-induced insulin secretion was significantly reduced in islets derived from FoxO1 βKO compared to wildtype (WT) islets (Fig. 1g,h). In vivo studies showed that the increase of insulin and decrease of glucose levels following GLB administration in WT mice were blunted in FoxO1 βKO (Fig. 1i,j).

### Cyb5r3 regulates GLB-induced insulin secretion

To understand the mechanism of impaired GLB-induced insulin secretion in FoxO1 βKO, we analyzed the role of its target Cyb5r3, which has been shown to affect glucose-induced insulin secretion (14). Cyb5r3 levels decreased in islets from FoxO1 βKO (Fig. 2a) and also following islet treatment with GLB *in vitro* for 96 h or in live mice for 4 weeks (Fig. 2b,c). To determine whether the impairment of GLB-induced insulin secretion caused by FoxO1 ablation is mediated by decreased expression of Cyb5r3, we restored Cyb5r3 in islets from FoxO1 βKO and analyzed GLB-induced insulin secretion. The reduced responsiveness to GLB in FoxO1 βKO islets was restored by adenovirus-mediated Cyb5r3 expression (Fig. 2d). These findings suggest that Cyb5r3 plays an important role in the regulation of GLB-induced insulin secretion by FoxO1.

**Fig. 2.**
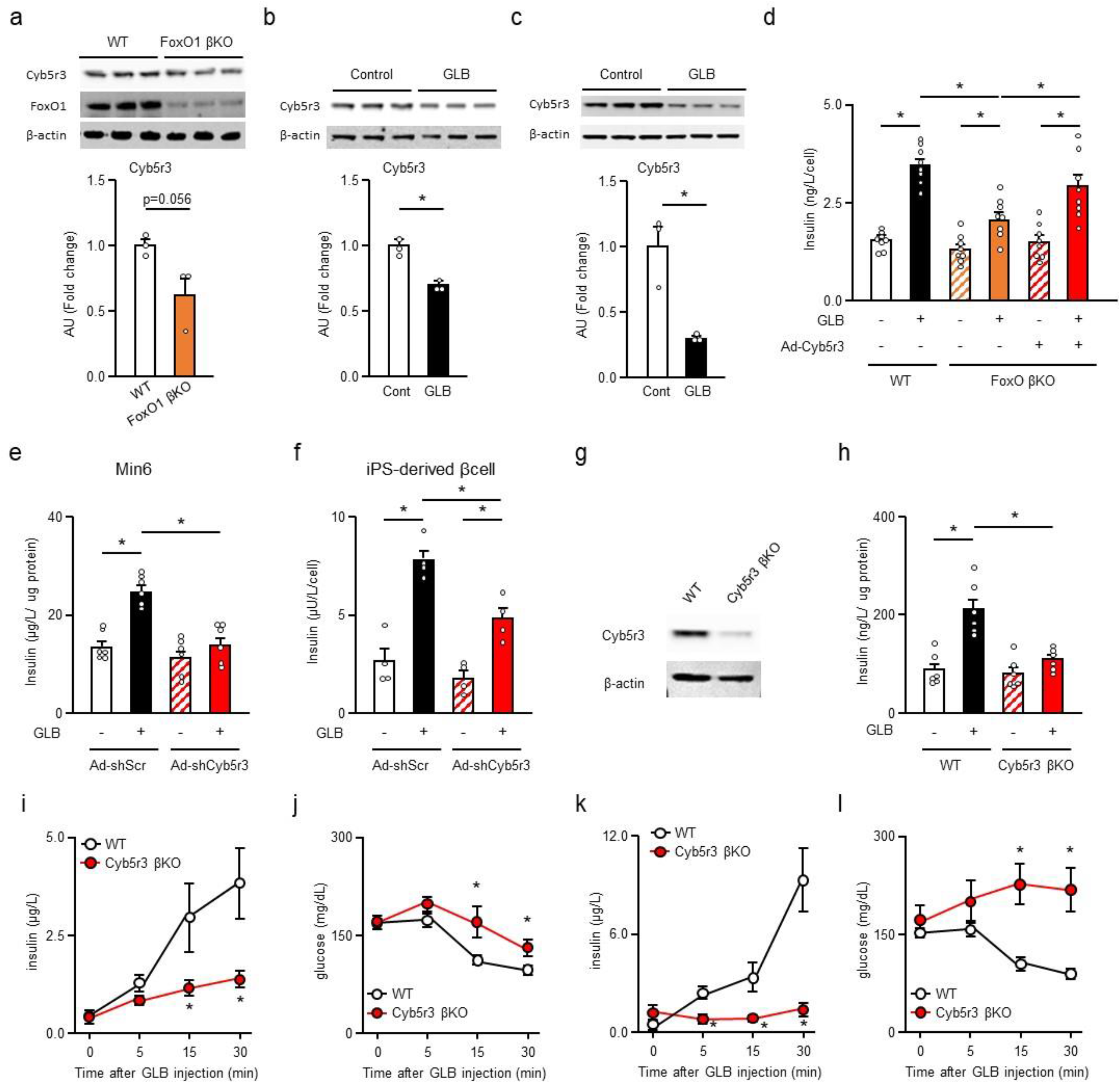
Cyb5r3 regulates GLB-induced insulin secretion. **a**, Cyb5r3 levels in islets from WT and FoxO1 βKO. **b**, Cyb5r3 levels in islets with or without GLB treatment for 96 h (n=3). c, Cyb5r3 levels in islets from mice with or without GLB treatment for 4 weeks (n=3). **d**, Adenovirus-mediated transduction of Cyb5r3 and GLB-induced insulin secretion in reaggregated islets from WT and FoxO1 βKO (n=8). **e,f**, Adenovirus-mediated Cyb5r3 knockdown and GLB-induced insulin secretion in MIN6 cells (n=6) (**e**) and human iPS-derived β-like-cells (n=4) (**f**). **g**, Cyb5r3 levels in WT and Cyb5r3 βKO islets. **h**, GLB-induced insulin secretion in islets from WT and Cyb5r3 βKO (n=6). **i,j**, Insulin (**i**) and glucose levels (j) after GLB administration in 10- to 20-week-old WT and Cyb5r3 βKO (n=7). **k,l**, Insulin (**k**) and glucose levels (**l**) after GLB administration in 40- to 50-week-old WT and Cyb5r3 βKO (n=4). \**P* < 0.05 by unpaired Student’s t-test, one-way ANOVA and two-way ANOVA followed by Tukey’s test. Data represent means ± SEM.

We then investigated the role of Cyb5r3 in GLB-induced insulin secretion. Knockdown of Cyb5r3 by adenoviral vector in MIN6 and human iPS cell-derived β-cells significantly reduced insulin secretion elicited by GLB (Fig. 2e,f). Cyb5r3 β-cell-specific knockout mice (Cyb5r3 βKO) islets were essentially unresponsive to GLB-induced insulin secretion compared to WT (Fig. 2g,h). Insulin secretion and hypoglycemic effect of GLB administration were also reduced in Cyb5r3 βKO compared to WT (Fig. 2i,j). The impairment was observed in mice of both genders (Fig. S2a,b) and was more pronounced in aging mice (40-50 weeks of age) (Fig. 2k,l). These results suggest that Cyb5r3 is in the FoxO1 pathway regulating GLB-induced insulin secretion.

### A glucose-sensing defect in Cyb5r3 **β**-cell knockouts

Next, we investigated the mechanism of impaired GLB-induced insulin secretion in Cyb5r3 βKO ^15^. We measured insulin and glucose levels in aging mice after administration of another third-generation SU, gliclazide (GLC), while refeeding after a 16-h fast, and during glucose or olive oil tolerance tests. Aging Cyb5r3 βKO hardly responded to GLC and showed modest changes in insulin and glucose levels. Differences with WT were more pronounced than in younger mice (Fig. 3a, Fig. S3a). Insulin levels in Cyb5r3 βKO tended to be lower one hour after refeeding but were higher at the end of the 16-h fast (Fig. 3b). Glucose levels prior to refeeding also trended higher in Cyb5r3 βKO, but the difference did not reach statistical significance (Fig. S3b). On the other hand, in glucose tolerance test, Cyb5r3 βKO showed a slow rise of insulin levels and marked glucose intolerance (Fig. 3c, Fig. S3c). In the olive oil tolerance test, insulin levels tended to be higher and 2-hr glucose levels lower in Cyb5r3 βKO (Fig. 3d, Fig. S3d). We saw similar findings in male and female young mice as well (Fig. S2c,d, Fig. S3e-j). In addition, Cyb5r3 βKO were able to maintain near-normal glucose levels under normal feeding and fasting conditions (Fig. S3a- d,f,h,j, time 0). These results show that Cyb5r3 βKO impairs the insulin response to SU and glucose, but not to secretagogues other than glucose, such as lipids and amino acids, suggesting that Cyb5r3 βKO has a specific defect in glucose sensing. Interestingly, this defect recapitulated the early abnormalities of insulin secretion seen in type 2 diabetes and pre-diabetes ^17–19^.

**Fig. 3.**
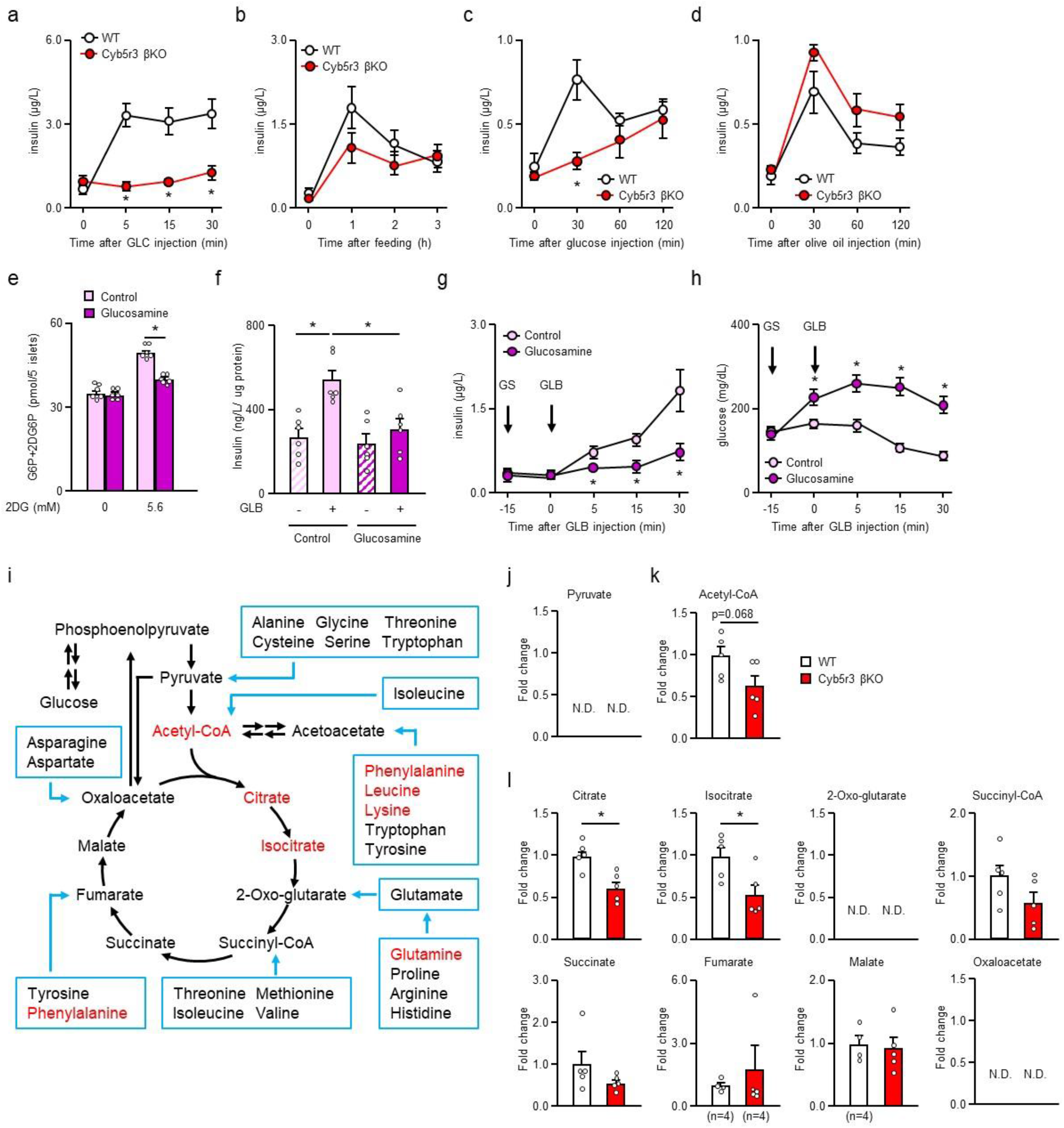
Cyb5r3 βKO affects glucose utilization and glucose responsiveness. **a-d**, Insulin levels after Gliclazide (GLC) (**a**), refeeding (**b**), glucose (**c**) and olive oil (**d**) administration in 40- to 50-week-old WT and Cyb5r3 βKO (n=4). **e**, Intracellular G6P+2DG6P content after 10-min 2DG treatment in islets preincubated with glucosamine (GS) for 1 h (n=6). **f**, GLB-induced insulin secretion in islets treated with or without GS for 1 h (n=6). **g,h**, Insulin (**g**) and glucose levels (h) in mice pretreated with or without GS 15 min prior to GLB injection (n=4). **i**, Schematic diagram of TCA cycle. Red color indicates metabolites depleted in Cyb5r3 βKO as determined by Metaboanalyst (P<0.11, VIP>1.28). **j-k**, Relative amounts of pyruvate (**j**), Acetyl-CoA (**k**) and TCA cycle metabolites (**l**) in WT and Cyb5r3 βKO in metabolomics analysis (n=5). \**P* < 0.05 by unpaired Student’s t-test, two-way ANOVA followed by Tukey’s test. Data represent means ± SEM.

### Reduced glycolysis associated with SU failure

Given these findings, we investigated whether glycolysis is involved in the failure of GLB-induced insulin secretion. To this end, we analyzed GLB-induced insulin secretion in the presence of glucosamine (GS), an inhibitor of the rate-limiting enzyme for glucose utilization in β-cells, Glucokinase (Gck) ^20^. Addition of 2- deoxyglucose (2DG) to islets increased the amount of intracellular 2DG6P (Fig. 3e). In contrast, the 2DG6P content of the GS pretreatment group showed a modest increase in intracellular 2DG6P content and was significantly lower than controls. At the same time, GLB-induced insulin secretion was suppressed in islets pretreated with GS (Fig. 3f). GS pretreatment of mice inhibited GLB induction of insulin secretion and resulted in significantly higher glucose levels than controls (Fig. 3g,h). These results suggest that GLB- induced insulin secretion requires a Gck-dependent step.

To understand the mechanism of this defect, we surveyed β-cell metabolism by metabolomics analysis. A total of 296 metabolites had a coefficient of variation (CV) ≤30% in quality control (QC) samples after removal of duplicate metabolites from both columns (Table S1). We imported the data set into SIMCA-p software and performed a supervised partial least square-discrimination analysis (PLS-DA). PLS-DA components effectively and distinctly separated Cyb5r3 βKO from WT islets (Fig. S4a), suggesting that Cyb5r3 deficiency caused significant changes in islet metabolite profile. Metaboanalyst-based pathway analysis yielded a list of enriched metabolites. The top 70 hits (P<0.11, VIP>1.28) demonstrated striking changes in amino acid and ketone body metabolism, as well as TCA cycle (Fig. S4b) (red letters in Fig. 3i). Analyses of individual metabolites indicated that pyruvate was undetectable in either genotype, probably as a result of its rapid conversion to acetyl-CoA (Fig. 3j). In Cyb5r3 βKO we observed a decrease in acetyl-CoA and in the early steps of the TCA cycle, citrate and isocitrate (Fig. 3k,l), while metabolites in later steps showed smaller or no differences between the two groups (Fig. 3l). These data indicate that metabolites in TCA cycle are reduced due to the lack of match between influx (anaplerotic flux) and outflow (CO2 production/energy and cataplerotic flux). Interestingly, amino acids that enter the TCA cycle via conversion to acetoacetate (leucine, lysine, phenylalanine, tyrosine) or glutamate (arginine, glutamine) were substantially decreased in Cyb5r3 βKO (Fig S4c,d). As most of the acetyl-CoA in β-cells is supplied from glucose, a possible explanation of these findings is that glycolysis is decreased in Cyb5r3 βKO, and the decrease in amino acids is a result of anapleurotic flux defects ^21^.

### Cyb5r3 binds Gck to regulate its levels and activity

To clarify whether Cyb5r3 is involved in glucose uptake, we analyzed 2-deoxyglucose (2DG) incorporation and Gck expression in islets from Cyb5r3 βKO. In WT islets, the amount of intracellular 2DG6P increased following addition of 2DG (Fig. 4a). In contrast, in Cyb5r3 βKO islets, intracellular 2DG6P hardly increased and total amount of endogenous G6P and 2DG6P derived from 2DG was significantly lower. Gck levels decreased substantially in Cyb5r3 βKO islets (Fig. 4b), while *Gck* mRNA increased somewhat and expression of other genes involved in SU-dependent insulin secretion was unchanged (Fig. 4c). To determine a causal relationship between Cyb5r3 and Gck levels, we performed loss- and gain-of-function experiments in MIN6 cells. Knockdown of Cyb5r3 decreased (Fig. 4d), while gain-of-function increased Gck protein levels (Fig. 4e). These results suggest that Cyb5r3 regulates Gck protein levels, not gene expression.

**Fig. 4.**
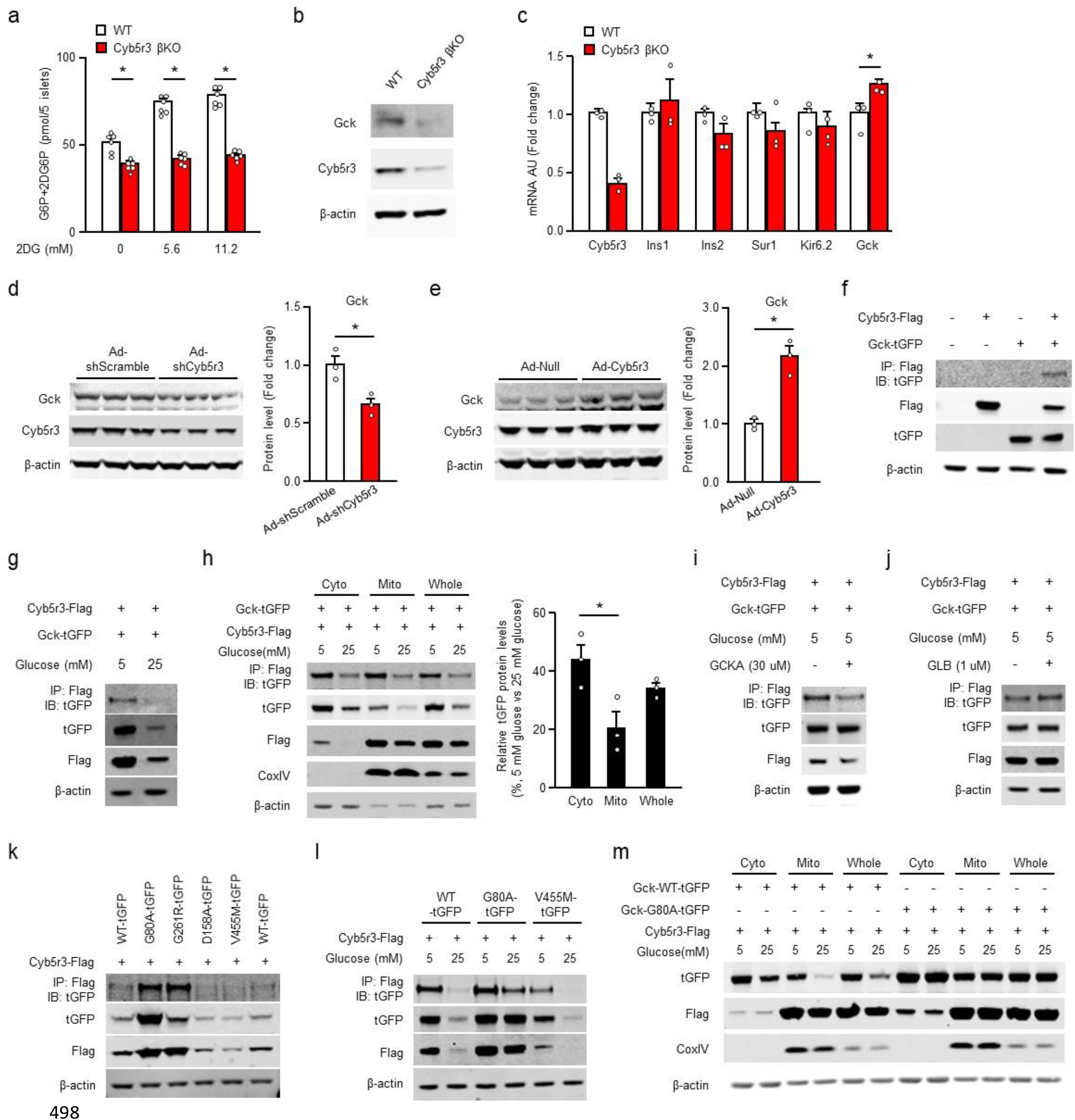
Cyb5r3 binds to Gck and regulates its protein level and activity. **a**, Intracellular G6P+2DG6P content after 10-min 2-deoxy-glucose (2DG) treatment in WT and Cyb5r3 βKO islets (n=6). **b**, Gck western blot in WT and Cyb5r3 βKO islets. **c**, mRNA expression in WT and Cyb5r3 βKO islets (n=3). **d,e**, Gck western blot in MIN6 cells following adenovirus-mediated knockdown (**d**) and gain-of-function of Cyb5r3 (**e**) (n=3). **f,g**, Immunoprecipitation (IP) and immunoblot (IB) of Cyb5r3 and Gck in HEK293 transfected with Flag-tagged Cyb5r3 and tGFP-tagged Gck (**f**) cultured for 6 h in 5 or 25 mM glucose (**g**). **h**, Western blot of subcellular fractions isolated from HEK293 expressing Flag-tagged Cyb5r3 and tGFP-tagged Gck cultured for 6 h in 5 or 25 mM glucose and relative tGFP levels in 25 mM glucose normalized by its levels in 5 mM glucose (n=3). Cyto: cytosol; mito: mitochondria. **i,j**, Western blot of HEK293 transfected with Flag-tagged Cyb5r3 and tGFP-tagged Gck and cultured for 1 h in 5 mM glucose with or without Gck activator (GCKA) (**i**) and GLB (**j**). **k**, Western blot of Cyb5r3 and WT or G80A, G261R, D158A and V455M mutant Gck in HEK293 cultured in 25 mM glucose medium. **l**, Western blot of Cyb5r3 and Gck in HEK293 transfected with Flag-tagged Cyb5r3 and tGFP-tagged WT, G80A or V455M mutant Gck cultured for 6 h in 5 or 25 mM glucose. **m**, Western blot of subcellular fractions from HEK293 transfected with Flag-tagged Cyb5r3 and tGFP-tagged WT or G80A Gck cultured for 3 h in 5 or 25 mM glucose. \**P* < 0.05 by unpaired Student’s t-test, one-way ANOVA followed by Tukey’s test. Data represent means ± SEM.

To determine whether Cyb5r3 acts directly on Gck, we examined protein-protein interactions using co-immunoprecipitation. In HEK293 cells co-expressing Flag-tagged Cyb5r3 and tGFP-tagged Gck, we detected Cyb5r3 in Gck immunoprecipitates and vice versa (Fig. 4f). Next, we analyzed the effects of glucose on the interaction. We observed binding of Cyb5r3 to Gck when we cultured cells for 6 h in 5 mM glucose; in contrast, following 6 h incubation at 25 mM glucose, a condition that increases Gck activity (Fig. 4g), Cyb5r3 binding was greatly reduced. Levels of Cyb5r3 and Gck decreased after incubation at 25 mM glucose for 6 h compared with 5 mM glucose. Both Gck and Cyb5r3 display a promiscuous subcellular localization ^15, 22–24^. In HEK293 cells co-expressing Cyb5r3 and Gck, Cyb5r3 mostly localized to mitochondria (Fig. 4h), while Gck localized to both cytoplasm and mitochondria. We performed subcellular fractionation to determine the site of the Cyb5r3/Gck interaction. We detected Cyb5r3 binding to Gck in both fractions, and both decreased following incubation in 25 mM glucose. Glucose-dependent Gck decrease was more pronounced in the mitochondrial than in the cytoplasmic fraction.

In addition, incubation with a Gck activator (GCKA) for 1 h in 5 mM glucose dissociated the Cyb5r3/Gck complex (Fig. 4i). In contrast, GLB stimulation for 1 h failed to dissociate the complex (Fig. 4j). As these data suggest that binding to Cyb5r3 modulates Gck activity, we asked whether Gck mutants with different activities differed in their binding to Cyb5r3.

Loss-of-function Gck mutations give rise to MODY2, a form of early-onset diabetes, whereas gain-of-function mutations result in persistent hypoglycemia of infancy ^25^. The critical test of our hypothesis was that loss-of-function mutants would be sequestered by Cyb5r3, whereas gain-of-function mutants would be Cyb5r3-independent. The findings bore out the hypothesis. Gck mutants G80A and G261R are the least active in β-cells, while V455M and D158A are the most active ^26^. In 25mM glucose, low-activity mutants G80A and G261R showed the strongest binding to Cyb5r3 (Fig. 4k). In contrast, high-activity mutants V455M and D158A showed the weakest binding to Cyb5r3. Compared to WT Gck, binding of G80A to Cyb5r3 was only modestly affected by raising glucose to 25mM, whereas V455M showed weak Cyb5r3 binding regardless of glucose concentrations (Fig. 4l). Gck and Cyb5r3 levels paralleled binding: G80A showed strong binding to Cyb5r3 and high Gck levels, whereas V455M showed weak binding to Cyb5r3 and low Gck levels. We wanted to determine whether changes in Gck levels were related to the subcellular localization of its binding to Cyb5r3. Under basal conditions (5mM glucose) cytoplasmic and mitochondrial localization of G80A was similar to WT (Fig. 4m). Protein content of G80A was unaffected by 25 mM glucose in any fraction. These results suggest that Cyb5r3 binding stabilizes Gck, while dissociation activates Gck and decreases its protein levels. Furthermore, as shown by the subcellular localization and binding to Cyb5r3 of G80A, Gck activity seems to be determined by Cyb5r3 binding, not by localization.

### Gck is required for GLB-induced insulin secretion in islets

To clarify whether the failure of GLB-induced insulin secretion in Cyb5r3 βKO is due to decreased Gck activity, we administered a Gck activator (GCKA) *in vitro* and *in vivo*. Treatment with GCKA restored GLB-induced insulin secretion in Cyb5r3 βKO islets (Fig. 5a). Insulin secretion was poorly responsive to GLB in Cyb5r3 βKO but improved dramatically following pre-administration of GCKA (Fig. 5b). Plasma insulin levels 30 min after GLB administration were significantly higher in mice pretreated with GCKA than controls (Fig. 5c). Pretreatment with GCKA also increased the glucose-lowering effect of GLB in Cyb5r3 βKO (Fig. 5d). Cyb5r3 βKO pretreated with GCKA showed significantly lower glucose levels 30 min after GLB administration (Fig. 5e). We also analyzed the effect of GCKA on GLB-induced insulin secretion in the *in vitro* GLB failure model. In islets treated with GLB for 96 h, not only Cyb5r3 but also Gck levels decreased (Fig. 2b,5f). Treatment with GCKA in islets with reduced Gck expression restored GLB-induced insulin secretion (Fig. 5g). Gck levels were also decreased in islets from mice treated with GLB for 4 weeks (Fig. 5h). GCKA treatment increased insulin and decreased glucose levels, reversing the impairment of GLB-induced insulin secretion (Fig. 5i,j). Stimulation with GLB for 96 h decreased FoxO1 and Cyb5r3 levels and tended to decrease Gck expression in human as well as murine islets (Fig. 5k). GLB- induced insulin secretion was also impaired in human islets treated with GLB for 96 h (Fig. 5l). Insulin secretion in human islets that developed impaired GLB-induced insulin secretion was restored by GCKA and GLB. These results suggest that the impairment of GLB-induced insulin secretion in Cyb5r3 βKO and the secondary failure of GLB treatment in human and murine islets is due to reduced Gck activity secondary to decreased Gck levels.

**Fig. 5.**
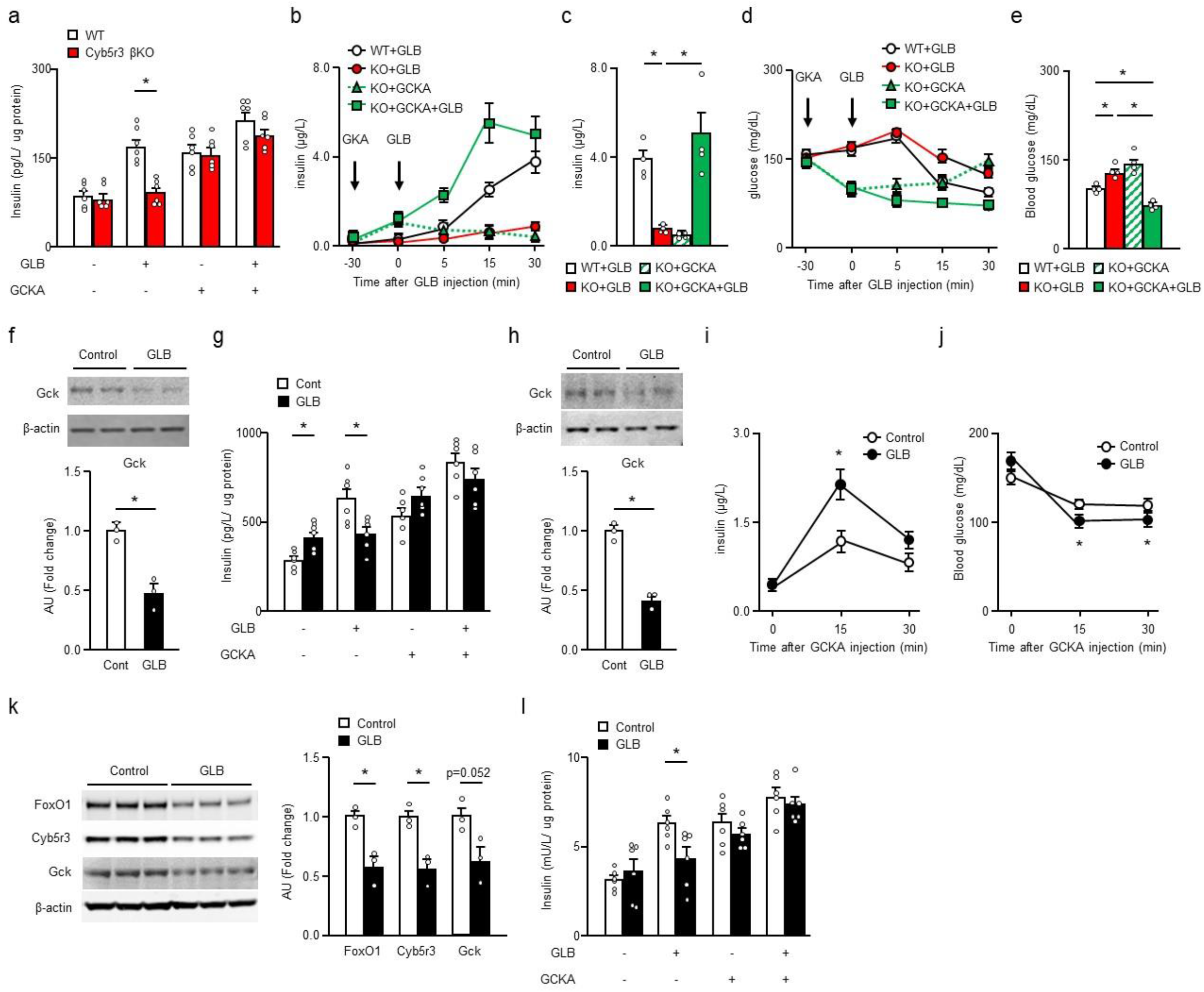
Gck activity is required for GLB-induced insulin secretion. **a**, GCKA and GLB-induced insulin secretion in islets from WT and Cyb5r3 βKO (n=6). **b-e**, Pre- administration of GCKA and insulin (**b**) or glucose (**d**) after GLB injection as well as 30 min insulin (**c**) and glucose (**e**) following GLB administration in WT and Cyb5r3 βKO (n=4). **f**, Gck western blot in islets cultured in with or without GLB for 96 h (n=3). **g**, GCKA and GLB-induced insulin secretion in islets treated with or without GLB for 96 h (n=6). **h**, Gck western blot in islets from mice with or without GLB treatment for 4 weeks (n=3). **i,j**, Insulin (**i**) and glucose levels (**j**) after GCKA injection in mice treated with GLB for 3 weeks (n=7). **k**, Western blot in human islets treated with or without GLB for 96 h (n=3). **l**, GCKA and GLB-induced insulin secretion in human islets treated with or without GLB for 96 h (n=6). \**P* < 0.05 by unpaired Student’s t-test, one-way ANOVA followed by Tukey’s test. Data represent means ± SEM.

### Cyb5r3 activator THII rescues secondary failure to GLB

Finally, we examined the effect of tetrahydroindenoindole (THII), a potential activator of Cyb5r3, on GLB- induced insulin secretion in the secondary failure model. The antioxidant THII protects against oxidative processes partly through Cyb5r3 ^27^. In fact, administration of THII from drinking water extends the lifespan of mice, and this effect is more pronounced in Cyb5r3 transgenic mice ^28^. While islets cultured with GLB alone for 4 days showed decreased GLB responsiveness to insulin secretion, THII treatment restored the GLB response to control levels (Fig. 6a). When administered *in vivo* with the diet, THII did not affect body weight (Fig. S5a). 2-week GLB administration lowered insulin levels, and addition of THII tended to increase them (Fig. S5b,c). Glucose levels tended to increase after 2-weeks GLB treatment, but were unchanged in the other groups (Fig. S5d). GLB-induced insulin secretion was impaired by chronic GLB treatment and was restored by co-administration of THII (Fig. 6b). Insulin levels 30 min after GLB administration were substantially decreased in the chronic GLB group and were restored by THII co-administration (Fig. 6c). Addition of THII to chronic GLB also restored the hypoglycemic effect of GLB (Fig. 6d,e). THII also rescued impaired GLB-induced insulin secretion in human islets cultured with GLB alone for 4 days (Fig. 6f). These data indicate that boosting Cyb5r3 activity may provide relief from SU secondary failure.

**Fig. 6.**
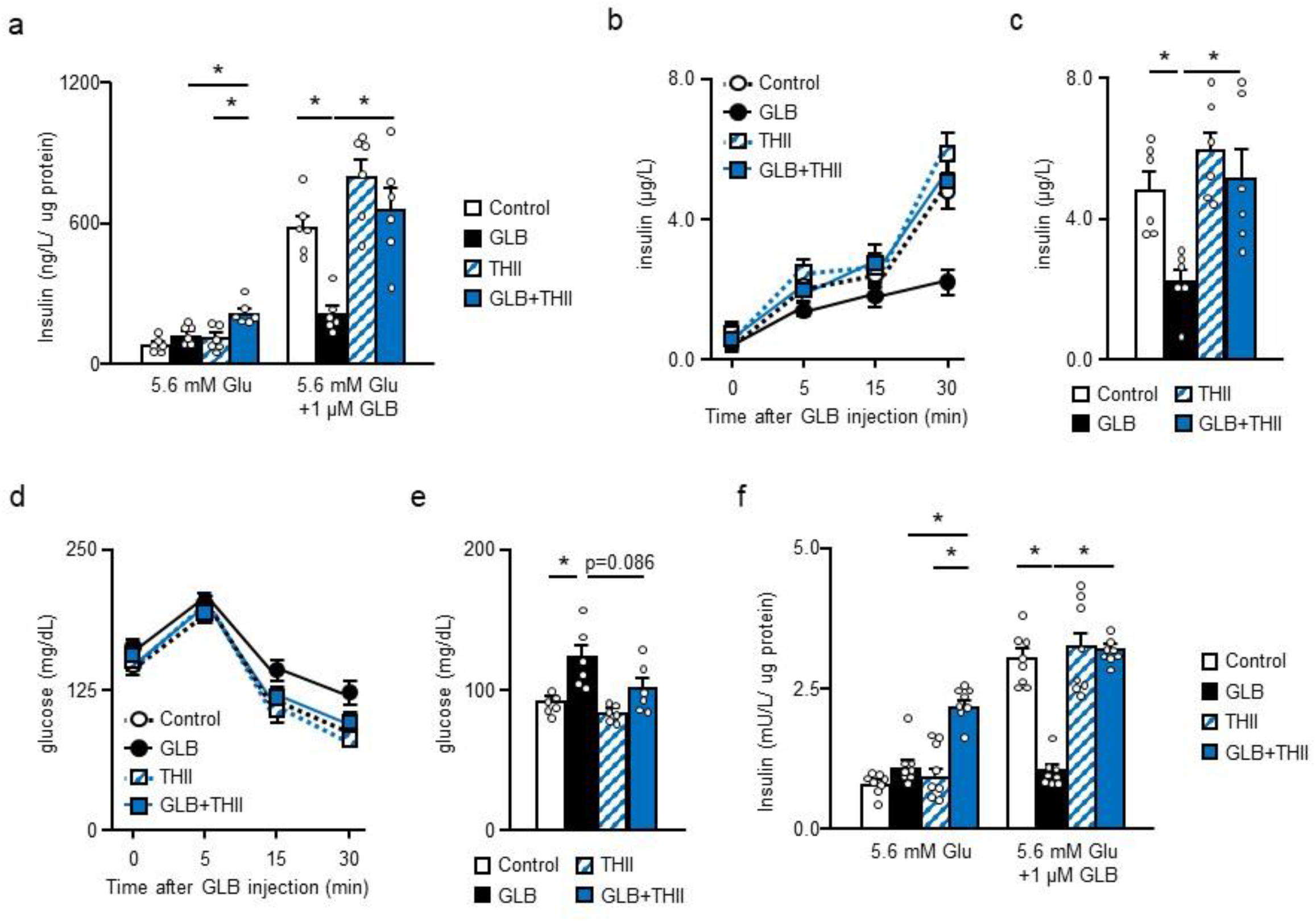
THII prevents GLB secondary failure. **a**, THII and GLB-induced insulin secretion in mouse islets treated with or without GLB for 96 h (n=6). **b,c**, Insulin levels after GLB injection (b) and at 30 min after GLB injection (c) after chronic GLB and THII treatment (n=6). **d,e**, Glucose levels after GLB injection (d) and at 30 min after GLB injection (e) after chronic GLB and THII treatment (n=6). **f**, THII and GLB-induced insulin secretion in human islets treated with or without GLB for 96 h (n=8). \**P* < 0.05 by two-way ANOVA followed by Tukey’s test. Data represent means ± SEM.

## Discussion

Key findings of this work are that (*i*) secondary failure to GLB administration is associated with decreased levels of FoxO1 and Cyb5r3; (*ii*) GLB-induced insulin secretion is natively impaired in FoxO1 βKO and Cyb5r3 βKO; (*iii*) Cyb5r3 regulates Gck levels and activity by direct binding; (*iv*) GCKA treatment restores GLB-induced insulin secretion in Cyb5r3 βKO and rescues secondary failure to GLB; (*v*) the Cyb5r3 activator, THII, rescues secondary failure to GLB; (*vi*) secondary failure to GLB is also associated decreased FoxO1, Cyb5r3 and Gck levels in human islets.

SU utilization had been declining for several years, owing to the introduction of Glp1 agonists and DPP4 inhibitors in the diabetes pharmacopeia. Why should we then care whether we can revamp it? For three main reasons: first, with time it has become apparent that Glp1 agonists and DPP4 inhibitors are not immune to secondary failures ^29–33^, and the former suffer from poor patient adherence ^34–36^. Second, as top- line findings from the comparative effectiveness GRADE study demonstrate, the five-year therapeutic failure rates of SU (72%) are comparable with those of other secretagogues or insulin (67 to 77%). Third, the GRADE study was solicited by the Center for Medicare and Medicaid Services as an independent assessment of cost-effectiveness of anti-diabetic treatments. SU cost on average 5% of Glp1 agonists, DPP4 inhibitors, and recombinant insulin ^37^. Considering runaway healthcare costs, being able to rely on older, cheaper drugs may offer a solution to this aspect of the diabetic care. Thus, restoring SU efficacy by preventing secondary failure has merit.

Studies of SU-dependent insulin secretion have focused on K_ATP_ channels and exchange protein directly activated by cAMP 2 (Epac2), a target of G protein-coupled receptors ^10^. But the mechanism of SU secondary failure remains unclear ^8^. Recently, the SUR1 binding site of GLB has been mapped ^11, 12^. Intracellular energetic states affect SUR1 structure and binding affinity of GLB to SUR1 ^11^. In a low-energy state characterized by increased Mg-ADP levels, GLB binds to the two nucleotide binding domains (NBD) of SUR1, causing them to dimerize and render the site inaccessible. We have previously shown that Cyb5r3 βKO is defective in ATP production, which may partly explain their unresponsiveness to SU ^15^. However, the present findings illustrate a more complex and pathophysiologically relevant picture. First, we found decreased Gck and impaired glucose uptake in vitro and following secondary failure to GLB, which were phenocopied by the Gck inhibitor, GS, and circumvented by the activator, GCKA, indicating that regulation of Gck is central to the mechanism by which Cyb5r3 ablation impairs the SU response. Consistent with our results, Gck knockout mice also show impaired GLB- and glucose-induced insulin secretion ^38^. These findings suggest that SU secondary failure involves a low energy state due to Gck dysfunction and reduced glucose utilization. The combined decrease of Cyb5r3 and Gck in β-cells can cause a low energy state and bring about a closed SUR1 structure, causing SU secondary failure.

Second, we provide a mechanism for this observation by showing that Cyb5r3 modulates Gck activity in β-cells. Gck regulatory protein (GKRP) and BAD are known to regulate Gck in β-cell and liver ^22, 39^. In the liver, GKRP binding stabilizes Gck, which is activated upon dissociation ^40, 41^. However, it’s controversial whether GKRP is expressed in β-cells. BAD binds to Gck on mitochondria and is involved in glucose-induced insulin secretion and apoptosis, but this interaction has not been assessed in SU failure ^24^. Cyb5r3 localizes to mitochondria and ER, and plays multiple roles in electron transfer and lipid synthesis^42^. We show that the bulk of the Cyb5r3/Gck complex localizes to mitochondria in HEK293 cells. However, whether there is subcellular specificity to the functional consequences of the Gck/Cyb5r3 interaction, similar to BAD-dependent cytochrome C release and apoptosis ^43^, remains to be seen. Apoptosis is unlikely to play a role in SU failure, since the latter can be rescued chemically without changes in cell number.

Regulation of the FoxO1/Cyb5r3 pathway bears intriguing similarities with incipient abnormalities of insulin secretion in diabetes that include a glucose-sensing defect and early-onset SU failure. Some clues as to the link of this early glucose-sensing defect with the more established steps of metabolic inflexibility and dedifferentiation associated with progression of β-cell failure and FoxO1 loss-of-function can be gleaned from the metabolomics analysis. The changes in the TCA cycle and amino acid metabolism pathways can be viewed as conducive to metabolic inflexibility ^13^ and may lead to dedifferentiation ^44^. In this regard, the demonstration that the Cyb5r3 activator, THII, can restore insulin secretion by reversing SU secondary failure provides a new target for the development of therapeutics that affect the FoxO1 pathway without targeting the transcription factor itself. More careful research into the pharmacology of Cyb5r3 will be required to turn these observations in to actionable development of diabetes therapeutics through elucidating the mechanism by which THII affects Cyb5r3.

## Acknowledgments

We thank Mr. Thomas Kolar and Ms. Ana Flete in Columbia University for exceptional technical support and members of the Accili laboratory for helpful discussions. This work was supported by the Manpei Suzuki Diabetes Foundation (HW) and NIH grants NIDDK 2R01DK64819 (DA), NIDDK 5P30DK63608 (Columbia Diabetes Research Center) and NIDDK P60DK020541 (Stable Isotope and Metabolomics Core Facility of the Albert Einstein College of Medicine).

## Author contributions

H.W. and D.A. designed the study and wrote the manuscript; H.W. performed all experiments; W. D., J.S., S.A., T. Kuo. and Y. M. performed experiments to characterize mouse models and islets; L.S. generated and performed iPS derived β-cells experiments. T. Kitamoto analyzed the data; I.K. analyzed metabolomics analysis; and R.C. provided THII and helpful suggestions.

## Declaration of interests

D.A. was a founder, director, and chair of the advisory board of Forkhead Therapeutics, Corp. This work is not related to the company.

## Resource availability

### Lead contact

Further information and requests for reagent and resource sharing should be directed to and will be fulfilled by the lead contact.

### Materials availability

All unique/stable reagents generated in this study are available from the lead contact with a completed Materials Transfer Agreement.

## Methods

### Animals

All animal experiments were in accordance with NIH guidelines, approved and overseen by the Columbia University Institutional Animal Care and Use Committee. All mice were housed in a climate-controlled room on a 12-h light/dark cycle with lights on at 7:00 AM and off at 7:00 PM and were fed standard chow (PicoLab rodent diet 20, 5053; Purina Mills). FoxO1-Venus, FoxO1 β-cell-specific knockout (FoxO1 βKO) and Cyb5r3 β-cell-specific knockout mice (Cyb5r3 βKO) have been described ^15, 16, 44^. C57BL/6 mice were from The Jackson Laboratory, Bar Harbor, ME. Mice were implanted a GLB-filled (20 mg/kg/day) (Sigma, St. Louis, MO) osmotic pump (ALZET, Cupertino, CA) intraperitoneally for 4 weeks. Treatment with THII during chronic GLB administration was performed as described ^28^. Mice were administered 100 µM THII in drinking water *ad libitum* for 4 weeks. We performed GLB (5 mg/kg), GLC (5 mg/kg) (Sigma), olive oil (10 ml/kg) (Sigma) tolerance tests in free feeding conditions and glucose (2 g/kg) and refeeding tolerance tests after a 16-h fast. GCKA (30 mg/kg) (Sigma) was injected intraperitoneally in Cyb5r3 βKO 30 min prior to GLB. Glucosamine (2g/kg) (Sigma) was injected intraperitoneally 15 min prior to GLB. Blood samples were obtained from the tail vein.

### Cell culture and isolated islet

MIN6 cells were cultured in Dulbecco’s modified Eagle medium (DMEM; ThermoFisher, Waltham, MA) containing 10% fetal bovine serum (FBS; Sigma) penicillin-streptomycin (ThermoFisher) at 37 °C and 5% CO2 and were used 48 h after adenovirus transduction. HEK 293 were cultured in DMEM with 10% FBS and transfected with plasmid for 24 h using Lipofectamine 3000 transfection reagent (ThermoFisher). HEK 293 were cultured in 5 or 25 mM glucose, 1 μM GLB and 30 μM GCKA for 1 to 6 h. β-like-cells from human induced pluripotent stem cells (iPSCs) were generated as described ^45^. As a quality control of differentiation, an aliquot of cells was stained with stage-specific markers of definitive endoderm, pancreatic progenitor, and β-cell differentiation stages. Islets were isolated from female mice by collagenase digestion as described ^13, 15^ and cultured in Roswell Park Memorial Institute (RPMI) 1640 medium (ThermoFisher) containing 10% FBS. For chronic GLB treatment, islets were cultured in RPMI 1640 containing 5.6 mM glucose, 1 μM GLB and 1 μM THII for 4 days. Human non-diabetic islets were obtained from the NIH’s Integrated Islet Distribution Program (IIDP) and cultured as described ^46^.

### Immunohistochemistry

Isolated islets were fixed in 4% PFA for 2 h after GLB treatment for 1 to 4 days. We used primary antibodies to GFP (A6455, 1:1000; ThermoFisher) and secondary anti-IgG antibodies conjugated with Alexa Fluor 488 (1:1,000; ThermoFisher).

### Adenovirus transduction

Adenoviruses encoding shCyb5r3 and Cyb5r3 have been described ^15^. β-like-cell clusters derived from iPSCs were dissociated using TrypLE Express (ThermoFisher) as described ^47^. Medium containing 2,000 cells and adenovirus encoding shCyb5r3 were placed in a round-bottom 96-well plate and cells were cultured for 2 days to reaggregate. Islets were dissociated into single cells using trypsin-EDTA (ThermoFisher) as described ^48^. Drops (20 µl of medium) containing 2,000 cells and adenovirus encoding Cyb5r3 were placed on top of a petri dish and cultured as hanging drops for 3 days. Pseudo-islets transduced with adenovirus encoding Cyb5r3 were used for insulin secretion assays.

### GLB-induced insulin secretion

Cells and islets were preincubated in KRBH buffer for 1 h in 5.6 mM glucose, followed by incubation in KRBH at 5.6 mM glucose with 1 μM GLB for 1 h at 37°C. At the end of the incubation, we collected islets by centrifugation and assayed supernatant and cell lysates for insulin. Insulin levels were normalized to protein content.

### Metabolites and biochemical analysis

Glucose was measured using a CONTOUR NEXT ONE (Ascensia Diabetes Care, Valhalla, NY), insulin with a Mouse (Mercodia,) or Human Insulin ELISA kit (both from Mercodia, Winston Salem, NC). In the 2-DG uptake assay, islets were preincubated in KRBH buffer for 1 h in 5.6 mM glucose, followed by incubation in KRBH in 0 to 11.2 mM 2-DG without glucose for 10 min at 37°C. Glucosamine (5 mM) was added to KRBH 10 min before replacing medium with KRBH containing 2-DG. 2-DG uptake was measured using a Glucose Uptake-Glo Assay kit (Promega, Madison, WI).

### Metabolomics analysis

Islets samples were extracted with 80% methanol with internal standards and freeze-thawed 3 times. After centrifugation the supernatant was transferred to a glass vial for LC injection. Samples were analyzed with ABsciex 6500+ with Ace PFP column and a iHILIC-p column (HILICON, Umeå, Sweden). A pooled QC sample was added to the sample list. This QC sample was injected six times for CV calculation for data quality control. We implemented the following CV parameters: metabolites with CVs < 20% were treated as accurate, CVs >20 and <30% as relatively accurate, CVs >30% as unreliable. Metabolites with a high CV in the QC samples were used only to perform further discovery. Data sets were analyzed using SIMCA-p.

### Western blotting and immunoprecipitation

Western blotting was performed as described ^41^. Islets and cultured cells were homogenized in ice cold CelLytic MT Cell Lysis Reagent (Sigma) with protease inhibitors. Mitochondrial and cytosol fractions were separated using a mitochondria isolation kit (ThermoFisher). Flag immunoprecipitation was performed with antibody conjugated-magnetic beads (OriGene, Rockville, MD). The following antibodies were obtained for immunoblotting: anti-FoxO1 (#2880, 1:1000), anti-Flag (#14793, 1:1000), anti-β-actin (#3770, 1:1000) (Cell Signaling Technology, Danvers, MA), anti-Gck (19666-1-AP, 1:500), anti-Cyb5r3 (10894-1-AP, 1:1000) (Proteintech, Chicago, IL), anti-tGFP (TA150075, 1:500; Origene), anti-CoxIV (ab33985, 1:500; Abcam, Cambridge, UK).

### Quantitative PCR

RNA was extracted from islets using RNeasy Mini Kit (Qiagen, Venlo, Netherlands). cDNA was synthesized by using qScript cDNA Synthesis Kit (QuantaBio, Beverly, MA) and PCR was performed with GoTaq qPCR Master Mix (Promega) with HPRT as the control gene. Primer sequences are available upon request.

### Statistical Analyses

Data are presented as mean ± SEM. Statistical analysis was performed with Student’s t-test for comparison between two groups, one-way ANOVA followed by Tukey’s test for multiple comparisons and two-way ANOVA followed by Tukey’s test for effects of two variables. Differences were considered significant at *P*< 0.05.

---

**Fig. S1.**
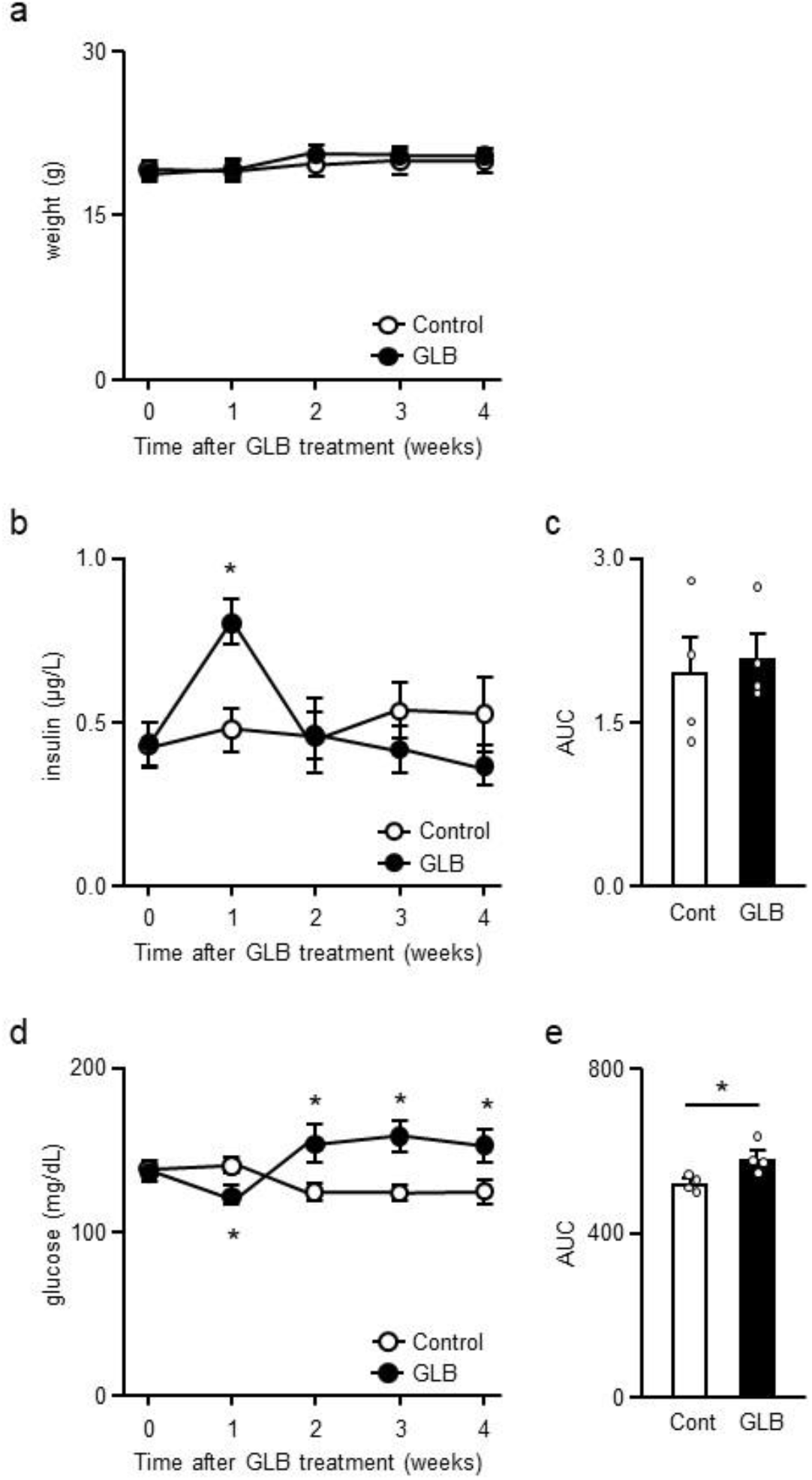
Change in insulin and glucose levels after GLB administration. **a**, Weight of mice treated with or without GLB for 4 weeks (n=4). **b,c**, Insulin (**b**) and area under curve (AUC) (**c**) in mice treated with or without GLB for 4 weeks (n=4). **d,e**, Glucose (**d**) and AUC (**e**) in mice treated with or without GLB for 4 weeks (n=4). \**P* < 0.05 by unpaired Student’s *t*-test. Data represent means ± SEM.

**Fig. S2.**
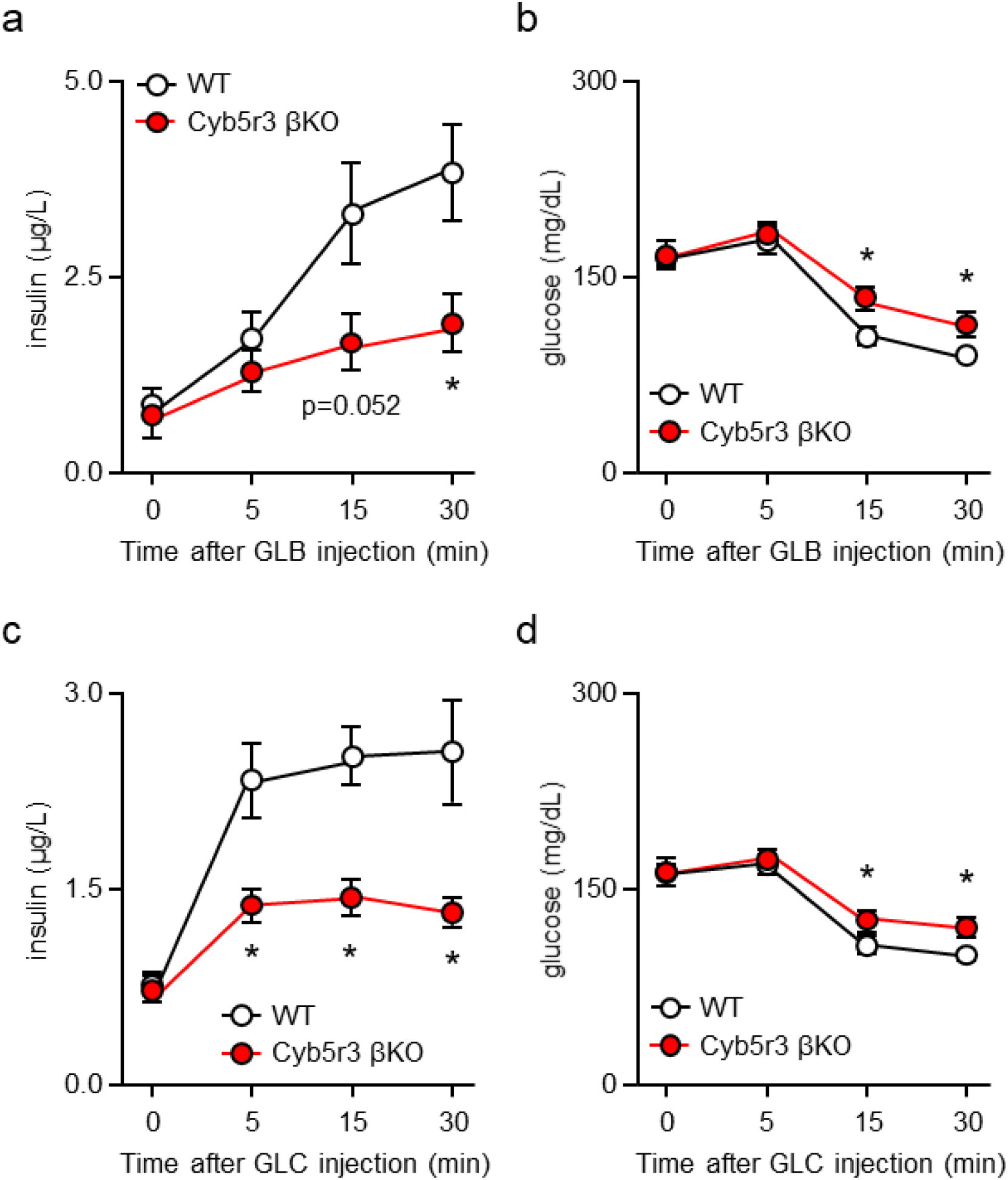
GLB-induced insulin secretion in male WT and Cyb5r3 βKO. **a,b**, Insulin (**a**) and glucose (**b**) after GLB administration in 10- to 20-week-old male WT and Cyb5r3 βKO (n=6). **c,d**, Insulin (**c**) and glucose (**d**) after GLC administration in 10- to 20-week-old male WT and Cyb5r3 βKO (n=6). \**P* < 0.05 by unpaired Student’s *t*-test. Data represent means ± SEM.

**Fig. S3.**
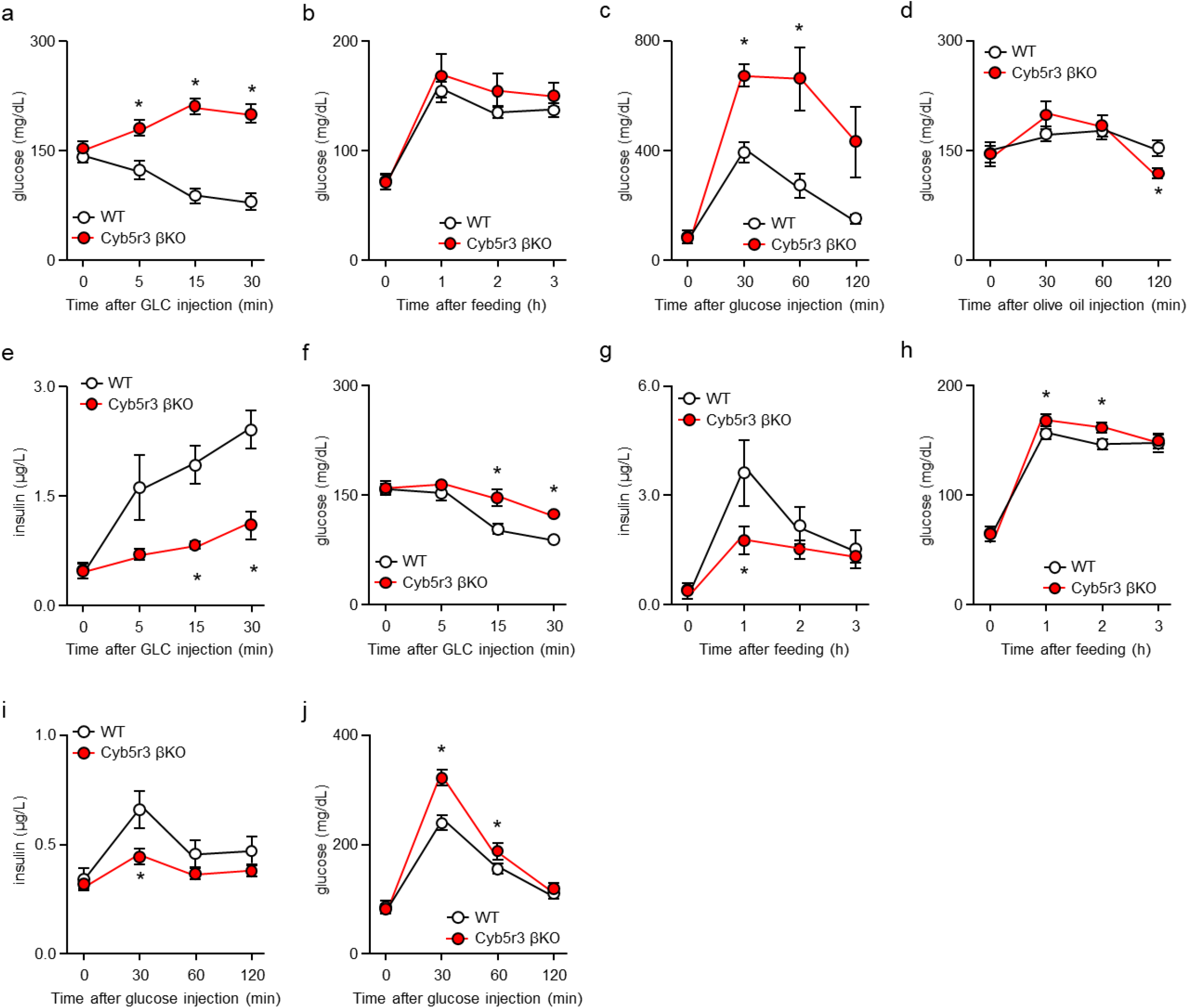
Insulin levels in WT and Cyb5r3 βKO. **a-d**, Glucose after GLC administration (**a**), refeeding (**b**), and glucose (**c**) or olive oil administration (**d**) in WT and Cyb5r3 βKO (40-50 weeks of age). **e-j**, Insulin (**e,g,i**) and glucose (**f,h,j**) after GLC administration (**e,f**), refeeding (**g,h**) and glucose administration (i**,j**) in WT and Cyb5r3 βKO (10-20 weeks of age) (n=5-7). \**P* < 0.05 by unpaired Student’s *t*-test. Data represent means ± SEM.

**Fig. S4.**
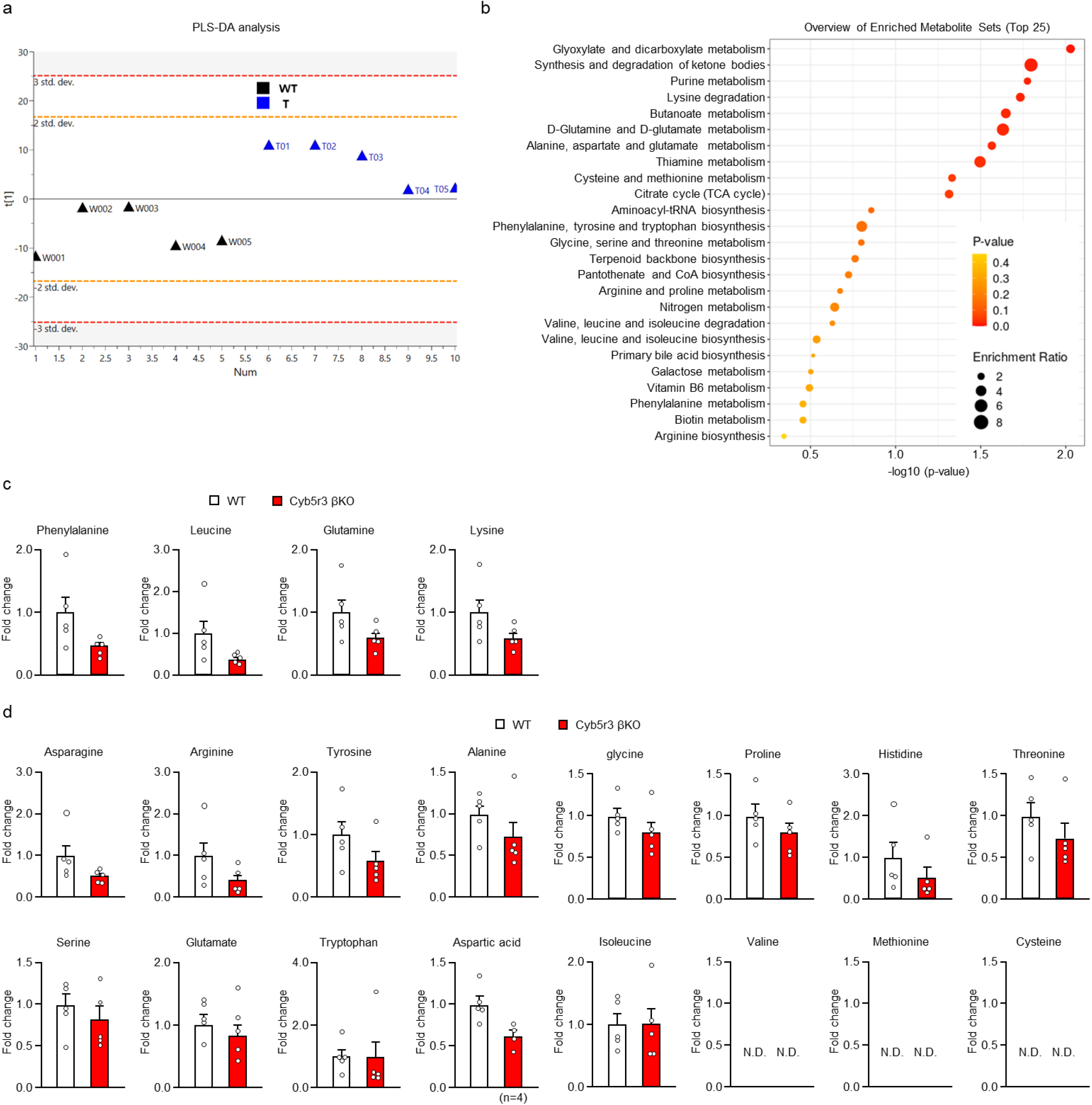
Metabolomics analysis and amino acid levels in islets from WT and Cyb5r3 βKO. **a**, Partial least square-discrimination (PLS-DA) analysis of metabolomic datasets in islets from WT and Cyb5r3 βKO. PLS-DA scores plots. The parameters for PLS-DA were 1 component with R2Xcum=0.255, R2Xcum=0.733, Q2cum=0.405). **b**, Overview of enriched metabolite sets (Top 25). Human Metabolite Database (HMDB) representation of top 70 hits (p<0.11, VIP value>1.28) used in enrichment analysis with Metaboanalyst. **c**, Relative amino acids levels in top 70 hits (p<0.11, VIP value>1.28) in islets form WT and Cyb5r3 βKO (n=5). **d**, Relative amino acid levels other than (**c**) in islets from WT and Cyb5r3 βKO (n=4-5). Data represent means ± SEM.

**Fig. S5.**
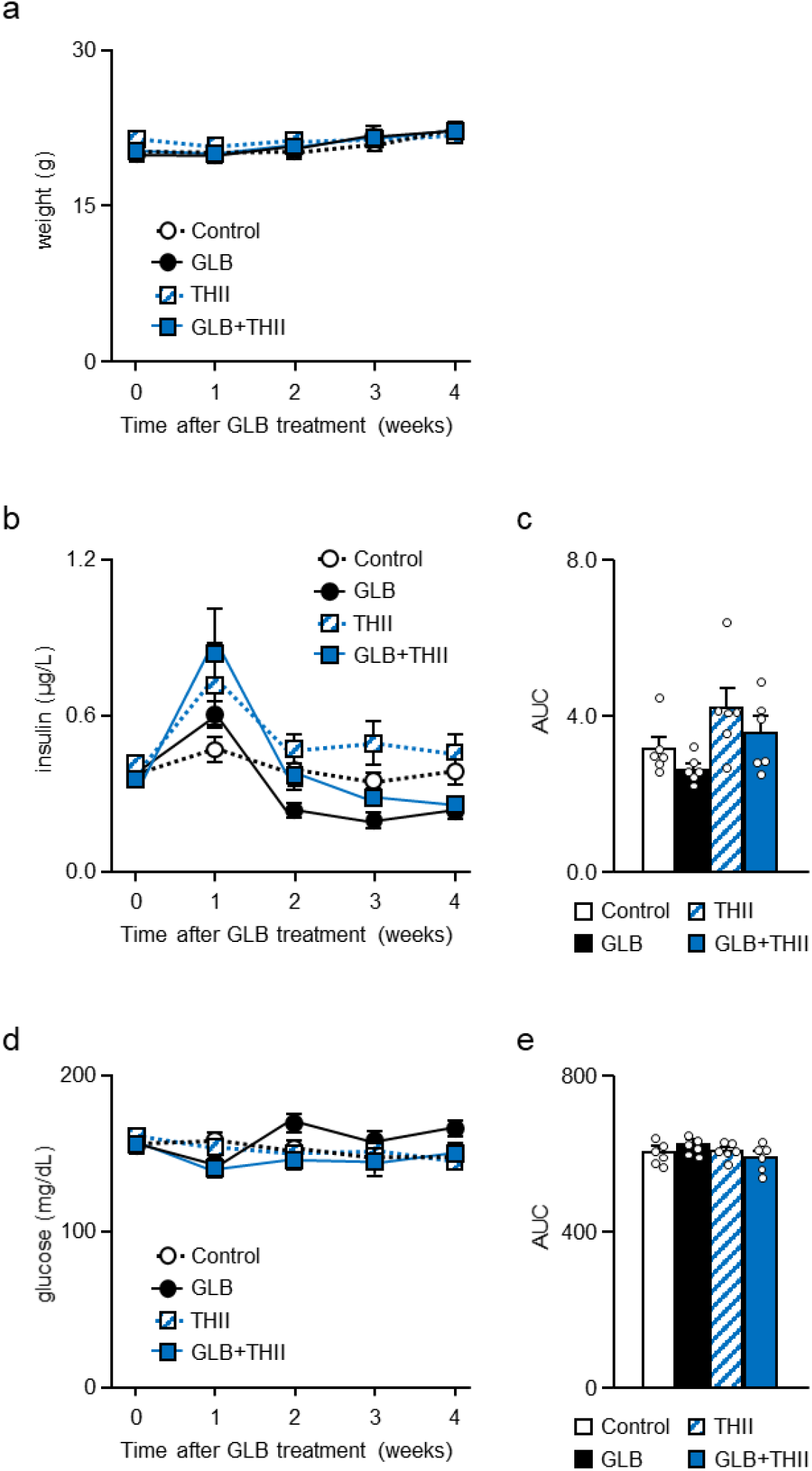
Effect of THII on weight, insulin and glucose. **a**, Weight of mice treated with GLB and THII for 4 weeks (n=6). **b,c**, Insulin (**b**) and area under curve (AUC) (**c**) in mice treated with GLB and THII for 4 weeks (n=6). **d,e**, Glucose (**d**) and AUC (**e**) in mice treated with GLB and THII for 4 weeks (n=6). Data represent means ± SEM.

