## Supplemental Table 1 for "Cyb5r3-based mechanism and reversal of secondary failure to sulfonylurea"

| Metabolite name | KO_01 | KO_02 | KO_03 | KO_04 | KO_05 | WT_01 | WT_02 | WT_03 | WT_04 | WT_05 | Average K | Average W | FC | P value | VIP value | CV |
|---|---|---|---|---|---|---|---|---|---|---|---|---|---|---|---|---|
| Pantothenic acid | 0.04543 | 0.0971 | 0.14125 | 0.1697 | 0.21168 | 0.26119 | 0.21471 | 0.28215 | 0.23579 | 0.25072 | 0.13303 | 0.24891 | 0.53445 | 0.00568 | 1.92257 | 21.249 |
| Inosine | 0.01835 | 0.01224 | 0.00649 | 0.01565 | 0.01966 | 0.02594 | 0.01967 | 0.02532 | 0.03375 | 0.03131 | 0.01448 | 0.0272 | 0.53233 | 0.00586 | 1.91841 | 13.1899 |
| Citric acid | 0.51828 | 0.34345 | 0.38597 | 0.67466 | 0.59976 | 0.80421 | 0.86992 | 0.61195 | 0.7881 | 0.9682 | 0.50443 | 0.80848 | 0.62392 | 0.00749 | 1.88503 | 1.31206 |
| adenine | 0.01228 | 0.00902 | 0.01429 | 0.0112 | 0.01943 | 0.02728 | 0.01955 | 0.02242 | 0.02504 | 0.01696 | 0.01324 | 0.02225 | 0.59525 | 0.00782 | 1.87887 | 8.06824 |
| Betaine | 0.09332 | 0.05363 | 0.0657 | 0.11569 | 0.09457 | 0.27528 | 0.17769 | 0.10636 | 0.20391 | 0.17046 | 0.08458 | 0.18674 | 0.45294 | 0.00853 | 1.86632 | 26.4128 |
| trans-4-Hydroxy-proline | 0.2895 | 0.19656 | 0.23209 | 0.23823 | 0.29136 | 0.45 | 0.31651 | 0.26919 | 0.41383 | 0.42321 | 0.24955 | 0.37455 | 0.66627 | 0.01275 | 1.80367 | 12.3825 |
| Cytidine 5'-diphosphocholine | 0.00839 | 0.00727 | 0.00594 | 0.01152 | 0.01198 | 0.01091 | 0.01475 | 0.01567 | 0.02286 | 0.01699 | 0.00902 | 0.01623 | 0.55553 | 0.01308 | 1.79946 | 2.75927 |
| N-Acetyl-DL-serine | 0.00029 | 0.00024 | 0.00031 | 0.00048 | 0.00035 | N/A | 0.00058 | 0.00047 | N/A | 0.00061 | 0.00033 | 0.00055 | 0.60493 | 0.01424 | 1.86923 | 3.52006 |
| Suberic acid | 0.00419 | 0.00443 | N/A | 0.00379 | 0.00418 | 0.01842 | 0.01227 | 0.00551 | 0.01772 | 0.0106 | 0.00415 | 0.0129 | 0.32153 | 0.01441 | 1.84155 | 19.0763 |
| Cystathionine | 0.01025 | 0.00268 | 0.00414 | 0.00583 | 0.0055 | 0.00864 | 0.01042 | 0.00836 | 0.01161 | 0.01239 | 0.00568 | 0.01028 | 0.55259 | 0.01535 | 1.7721 | 13.5475 |
| 2,3-Diaminopropionic acid | 0.01576 | 0.00817 | 0.01671 | 0.01281 | 0.01433 | 0.01641 | 0.02099 | 0.01671 | 0.02037 | 0.02145 | 0.01356 | 0.01918 | 0.70676 | 0.01611 | 1.76366 | 7.77825 |
| N-Acetyl-L-phenylalanine | 0.00117 | 0.00067 | 0.00069 | 0.00136 | 0.00043 | 0.00273 | 0.00202 | 0.00071 | 0.00224 | 0.00277 | 0.00086 | 0.00209 | 0.41193 | 0.01748 | 1.74901 | 22.5493 |
| Cytidine 2':3'-cyclic monophosphate | 3.6E-05 | 1.2E-05 | 1.6E-05 | 6.3E-05 | 2.3E-05 | 0.00023 | 0.0001 | N/A | 0.00053 | 0.00024 | 3E-05 | 0.00027 | 0.10982 | 0.01778 | 1.80566 | 9.82055 |
| Thiamine monophosphate | 0.00292 | 0.00253 | 0.00357 | 0.00283 | 0.0039 | 0.00527 | 0.00335 | 0.00571 | 0.00388 | 0.006 | 0.00315 | 0.00484 | 0.65112 | 0.01937 | 1.73002 | 26.1918 |
| 2-Hydroxyglutaric acid | 0.00603 | 0.0046 | 0.00808 | 0.00493 | 0.00777 | 0.01975 | 0.01841 | 0.00353 | 0.0216 | 0.01457 | 0.00628 | 0.01557 | 0.4034 | 0.02272 | 1.69938 | 25.8874 |
| taurochenodeoxycholate | 0.13868 | 0.12748 | 0.14443 | 0.22188 | 0.19999 | 0.31201 | 0.17939 | 0.21222 | 0.29574 | 0.27342 | 0.16649 | 0.25456 | 0.65405 | 0.02316 | 1.69558 | 8.5853 |
| trans-Cinnamic acid | 0.89767 | 1.01523 | 1.17346 | 0.84495 | 1.02902 | 1.27642 | 1.09873 | 1.33088 | 1.05194 | 1.27794 | 0.99207 | 1.20718 | 0.8218 | 0.02679 | 1.66601 | 11.3664 |
| Cytidine 5'-monophosphate | 0.04312 | 0.02639 | 0.03747 | 0.05008 | 0.04421 | 0.05287 | 0.04726 | 0.0478 | 0.05753 | 0.06312 | 0.04026 | 0.05371 | 0.74943 | 0.02748 | 1.66076 | 11.6628 |
| cis-4,7,10,13,16,19-Docosahexaenoic acid | 1.0173 | 0.74493 | 1.01079 | 1.09448 | 1.01447 | 1.0618 | 1.34966 | 1.08016 | 1.31931 | 1.20038 | 0.97639 | 1.20226 | 0.81213 | 0.02786 | 1.65787 | 12.1427 |
| 3-Hydroxy-3-methylglutaric acid | 0.00197 | 0.00131 | 0.00127 | 0.0011 | 0.00147 | 0.00498 | 0.00299 | 0.00175 | 0.00302 | 0.00212 | 0.00142 | 0.00297 | 0.4785 | 0.02787 | 1.65776 | 9.05366 |
| Isocitric acid | 0.11329 | 0.10864 | 0.11875 | 0.2129 | 0.31557 | 0.25103 | 0.37847 | 0.2142 | 0.35804 | 0.40264 | 0.17383 | 0.32087 | 0.54174 | 0.02793 | 1.65736 | 16.5284 |
| N-acetyl-histidine | 0.00383 | 0.00215 | 0.00175 | 0.0087 | 0.00263 | 0.01401 | 0.01009 | 0.00229 | 0.01509 | 0.01138 | 0.00381 | 0.01057 | 0.36056 | 0.03109 | 1.63439 | 9.68316 |
| Adenylosuccinic acid | 0.02258 | 0.01368 | 0.02915 | 0.03913 | 0.02417 | 0.03963 | 0.03275 | 0.0392 | 0.04665 | 0.03349 | 0.02574 | 0.03835 | 0.67124 | 0.03228 | 1.62619 | 16.6648 |
| glycochenodeoxycholate | 0.01405 | 0.00924 | 0.00618 | 0.02059 | 0.01546 | 0.02726 | 0.0149 | 0.01846 | 0.0271 | 0.02232 | 0.0131 | 0.02201 | 0.59542 | 0.03366 | 1.61693 | 3.57189 |
| taurohyodeoxycholate | 0.04217 | 0.03379 | 0.03796 | 0.07386 | 0.0559 | 0.09203 | 0.05499 | 0.06539 | 0.09247 | 0.07146 | 0.04874 | 0.07527 | 0.64751 | 0.03418 | 1.61345 | 6.81186 |
| glycoursodeoxycholate | 0.01301 | 0.0098 | 0.00689 | 0.02056 | 0.01525 | 0.02667 | 0.01479 | 0.01701 | 0.02697 | 0.02289 | 0.0131 | 0.02167 | 0.60468 | 0.03655 | 1.59829 | 3.59838 |
| glycodeoxycholate | 0.01365 | 0.00986 | 0.00574 | 0.01971 | 0.01519 | 0.02608 | 0.01466 | 0.01681 | 0.02822 | 0.022 | 0.01283 | 0.02155 | 0.59535 | 0.03826 | 1.58769 | 4.18789 |
| Guanosine monophosphate | 0.0002 | 0.00027 | 0.00019 | 0.00074 | 0.0006 | 0.00058 | 0.00065 | 0.00093 | 0.00065 | 0.00078 | 0.0004 | 0.00072 | 0.55757 | 0.03856 | 1.5859 | 18.7625 |
| taurodeoxycholate | 0.04377 | 0.03917 | 0.04168 | 0.08256 | 0.06193 | 0.10633 | 0.05897 | 0.06926 | 0.10614 | 0.08372 | 0.05382 | 0.08488 | 0.6341 | 0.03921 | 1.58199 | 5.08681 |
| glycocholate | 0.00323 | 0.00309 | 0.00279 | 0.00423 | 0.00351 | 0.0065 | 0.00365 | 0.0034 | 0.00589 | 0.00534 | 0.00337 | 0.00496 | 0.68003 | 0.04302 | 1.5599 | 5.48949 |
| glycohyodeoxycholate | 0.00935 | 0.00742 | 0.00504 | 0.01713 | 0.01212 | 0.02294 | 0.0117 | 0.01394 | 0.02214 | 0.01663 | 0.01021 | 0.01747 | 0.5846 | 0.04415 | 1.5536 | 4.97315 |
| Niacinamide | 0.19156 | 0.13034 | 0.19127 | 0.1295 | 0.18092 | 0.32591 | 0.19682 | 0.23004 | 0.46561 | 0.21206 | 0.16472 | 0.28609 | 0.57577 | 0.04857 | 1.53 | 5.45716 |
| D-Galactose | 0.0023 | 0.00224 | 0.00176 | 0.00166 | 0.00154 | 0.0013 | 0.00151 | 0.00054 | 0.00147 | 0.00171 | 0.0019 | 0.00131 | 1.45346 | 0.04886 | 1.52845 | 17.7403 |
| cytosine | 0.05726 | 0.02979 | 0.03203 | 0.05267 | 0.04418 | 0.0627 | 0.05395 | 0.04386 | 0.07438 | 0.07379 | 0.04318 | 0.06174 | 0.69951 | 0.04895 | 1.52802 | 8.8123 |
| 4-aminobutyrate | 0.30245 | 0.12945 | 0.34402 | 0.41663 | 0.30708 | 0.44178 | 0.39533 | 0.33892 | 0.45754 | 0.58294 | 0.29993 | 0.4433 | 0.67657 | 0.05032 | 1.52104 | 20.5678 |
| DL-3-Hydroxy-3-methylglutaryl coenzyme A | 7.9E-05 | 7.1E-05 | 7.7E-06 | 8.3E-05 | 4.5E-05 | 8.6E-05 | 4.6E-05 | 0.00018 | 0.00016 | 0.00024 | 5.7E-05 | 0.00014 | 0.40183 | 0.05096 | 1.5178 | 22.8112 |
| Creatine | 0.63767 | 0.31123 | 0.16647 | 0.356 | 0.32947 | 0.74581 | 0.50346 | 0.42226 | 0.66891 | 0.55337 | 0.36017 | 0.57876 | 0.62231 | 0.05252 | 1.51006 | 5.10774 |
| N-Methyl-glutamic acid | 0.01117 | 0.00937 | 0.00828 | 0.02842 | 0.01493 | 0.03661 | 0.02003 | 0.01214 | 0.04882 | 0.0472 | 0.01443 | 0.03296 | 0.43794 | 0.05332 | 1.50615 | 9.44347 |
| Isovaleraldehyde | 0.00136 | 0.00092 | 0.00241 | 0.00208 | 0.00123 | 0.00214 | 0.00196 | 0.00228 | 0.00268 | 0.00234 | 0.0016 | 0.00228 | 0.70186 | 0.05456 | 1.50014 | 16.1295 |
| Betaine | 0.27865 | 0.24222 | 0.18227 | 0.28928 | 0.26895 | 0.8166 | 0.3358 | 0.31425 | 0.39762 | 0.44649 | 0.25227 | 0.46215 | 0.54587 | 0.05523 | 1.49692 | 8.74721 |
| Pyridoxal 5'-phosphate | 0.00184 | 0.00264 | 0.00114 | 0.00738 | 0.01288 | 0.0113 | 0.00599 | 0.01208 | 0.01083 | 0.01344 | 0.00518 | 0.01073 | 0.48268 | 0.06116 | 1.46969 | 19.1562 |
| N-Acetyl-L-leucine | 0.00092 | 0.00125 | 0.00078 | 0.00087 | 0.00169 | 0.00431 | 0.00183 | 0.00121 | 0.00218 | 0.00192 | 0.0011 | 0.00229 | 0.48086 | 0.06455 | 1.45487 | 13.4893 |
| 2,5-Dimethylpyrazine | 9.9E-05 | 0.00075 | 0.00084 | 0.00017 | 0.00042 | 0.00121 | 0.00092 | 0.00075 | 0.00047 | 0.001 | 0.00046 | 0.00087 | 0.52549 | 0.067 | 1.44446 | 28.9758 |
| Acetyl CoA | 0.00344 | 0.00335 | 0.00205 | 0.00656 | 0.00645 | 0.00538 | 0.00487 | 0.00901 | 0.00712 | 0.00821 | 0.00437 | 0.00692 | 0.63194 | 0.06795 | 1.44051 | 10.2762 |
| Butyryl-carnitine | 0.00796 | 0.00651 | 0.00492 | 0.00905 | 0.01026 | 0.01875 | 0.01055 | 0.00705 | 0.01621 | 0.01058 | 0.00774 | 0.01263 | 0.61278 | 0.0681 | 1.43987 | 2.127 |
| Propionyl Carnitine | 0.00533 | 0.00973 | 0.00853 | 0.01175 | 0.01237 | 0.02092 | 0.01406 | 0.01226 | 0.01137 | 0.01163 | 0.00954 | 0.01405 | 0.67924 | 0.07263 | 1.4215 | 13.7201 |
| 5-Hydroxymethyluracil | 0.00312 | 0.00063 | 0.00165 | 0.00061 | 0.00338 | 0.00974 | 0.00316 | 0.00294 | 0.00626 | 0.00266 | 0.00188 | 0.00495 | 0.37958 | 0.07267 | 1.42134 | 18.1984 |
| Nonadecanoic acid | 0.16226 | 0.18904 | 0.24028 | 0.20247 | 0.19226 | 0.23452 | 0.20498 | 0.22236 | 0.22436 | 0.24522 | 0.19726 | 0.22629 | 0.87172 | 0.07703 | 1.40435 | 8.2334 |
| Phenylalanine | 0.45134 | 0.26239 | 0.20361 | 0.39254 | 0.45937 | 1.49261 | 0.55644 | 0.34 | 0.84171 | 0.59072 | 0.35385 | 0.76429 | 0.46298 | 0.0806 | 1.3909 | 24.3839 |
| 5-Methylcytosine | 0.00262 | 0.00242 | 0.00118 | 0.0033 | 0.00151 | 0.0029 | 0.00259 | 0.00351 | 0.0031 | 0.00306 | 0.00221 | 0.00303 | 0.72848 | 0.08119 | 1.38872 | 19.4126 |
| Guanosine 3′,5′-cyclic monophosphate | 0.02531 | 0.05776 | 0.03633 | 0.19828 | 0.16153 | 0.10411 | 0.1954 | 0.16326 | 0.22 | 0.20056 | 0.09584 | 0.17667 | 0.5425 | 0.08188 | 1.38616 | 18.286 |
| Glycolate | 2.9E-05 | 3.2E-05 | 8.2E-05 | 8E-05 | 4.3E-06 | 0.00015 | 0.00014 | 1.2E-05 | 0.00011 | 1E-04 | 4.5E-05 | 0.0001 | 0.4428 | 0.08309 | 1.38175 | 18.9185 |
| 5'-Deoxy-5'-(methylthio)adenosine | 0.01976 | 0.01648 | 0.0201 | 0.03315 | 0.04001 | 0.02508 | 0.03832 | 0.03418 | 0.04615 | 0.04376 | 0.0259 | 0.0375 | 0.69072 | 0.08412 | 1.37798 | 5.67017 |
| N,N,N-Trimethyllysine | 0.03797 | 0.0122 | 0.01594 | 0.0198 | 0.02434 | 0.05011 | 0.0275 | 0.01963 | 0.04907 | 0.03655 | 0.02205 | 0.03657 | 0.60301 | 0.0868 | 1.36841 | 11.495 |
| alpha-Methylhistidine | 8.9E-05 | 0.00044 | 0.00063 | 0.00037 | 0.00042 | 0.00193 | 0.00039 | 0.0006 | 0.00058 | 0.00161 | 0.00039 | 0.00102 | 0.38165 | 0.08682 | 1.36835 | 25.6103 |
| Leucine | 1.08908 | 0.57315 | 0.64516 | 0.87844 | 1.25841 | 5.21599 | 1.56734 | 0.80052 | 2.51664 | 1.78403 | 0.88885 | 2.37691 | 0.37395 | 0.08993 | 1.35746 | 23.3142 |
| Inosine 5'-monophosphate | 0.04945 | 0.07439 | 0.0768 | 0.10462 | 0.09992 | 0.10002 | 0.10512 | 0.14194 | 0.08917 | 0.09862 | 0.08104 | 0.10698 | 0.75753 | 0.09051 | 1.35543 | 24.4101 |
| Carnitine (Carnitine_C0) | 0.25184 | 0.18139 | 0.18058 | 0.22141 | 0.24444 | 0.26554 | 0.22902 | 0.22388 | 0.26559 | 0.26673 | 0.21593 | 0.25015 | 0.8632 | 0.09342 | 1.3455 | 4.42613 |
| S-Adenosylmethionine | 0.23905 | 0.10403 | 0.14995 | 0.72466 | 0.92701 | 1.24365 | 0.60775 | 0.33708 | 1.51768 | 0.99454 | 0.42894 | 0.94014 | 0.45625 | 0.09492 | 1.34047 | 11.3949 |
| Thiamine | 0.00205 | 0.00041 | 0.00089 | 0.00064 | 0.00147 | 0.0029 | 0.00112 | 0.00158 | 0.00183 | 0.00197 | 0.00109 | 0.00188 | 0.58019 | 0.09575 | 1.3377 | 19.8677 |
| Kynurenic acid | 0.00732 | 0.01788 | 0.02247 | 0.01792 | 0.01413 | 0.0216 | 0.02088 | 0.02197 | 0.02063 | 0.01879 | 0.01594 | 0.02077 | 0.7675 | 0.09916 | 1.32649 | 18.7983 |
| Trimethylamine | 0.00874 | 0.00473 | 0.00533 | 0.00862 | 0.00792 | 0.00787 | 0.00904 | 0.00734 | 0.01081 | 0.01008 | 0.00707 | 0.00903 | 0.78283 | 0.10417 | 1.31042 | 4.96178 |
| DL-Homocysteine | 5E-06 | 9.9E-06 | 1E-05 | 6E-06 | 2.6E-06 | 5.6E-06 | 7.6E-06 | 4.9E-05 | 3.9E-05 | 1.3E-05 | 6.8E-06 | 2.3E-05 | 0.29618 | 0.10985 | 1.2928 | 29.4525 |
| Glutamine | 0.44127 | 0.16905 | 0.27518 | 0.27654 | 0.35443 | 0.88595 | 0.38742 | 0.27023 | 0.57855 | 0.42861 | 0.30329 | 0.51015 | 0.59452 | 0.11082 | 1.28986 | 4.49047 |
| 2-Isopropylmalic acid | 0.00189 | 0.0039 | 0.00419 | 0.00557 | 0.00476 | 0.00509 | 0.00558 | 0.00513 | 0.00541 | 0.00473 | 0.00406 | 0.00519 | 0.78314 | 0.11248 | 1.28485 | 14.6493 |
| Lysine | 0.51241 | 0.21807 | 0.31146 | 0.31499 | 0.4251 | 1.06481 | 0.46789 | 0.31372 | 0.66326 | 0.49771 | 0.35641 | 0.60148 | 0.59255 | 0.11405 | 1.28016 | 4.04047 |
| Adenosine 3',5'-diphosphate | 0.00651 | 0.00809 | 0.00365 | 0.0206 | 0.02506 | 0.01722 | 0.01214 | 0.03665 | 0.02218 | 0.02936 | 0.01278 | 0.02351 | 0.54368 | 0.11458 | 1.27859 | 20.8302 |
| Pimelic acid | 0.68326 | 0.72368 | 0.93615 | 0.83074 | 0.70683 | 0.86386 | 0.80955 | 0.91065 | 0.83385 | 0.91895 | 0.77614 | 0.86737 | 0.89481 | 0.11655 | 1.27276 | 5.81408 |
| Choline | 1.84663 | 1.1941 | 1.67551 | 2.49679 | 2.14778 | 3.53821 | 2.12594 | 1.5525 | 2.85828 | 2.81822 | 1.87216 | 2.57863 | 0.72603 | 0.1192 | 1.26502 | 13.9571 |
| glycolithocholate | 0.0006 | 0.00069 | 0.00013 | 0.00111 | 0.00084 | 0.00138 | 0.00051 | 0.00099 | 0.00119 | 0.00122 | 0.00067 | 0.00106 | 0.63821 | 0.12037 | 1.26165 | 7.17725 |
| 3-Methylhistidine | 0.00713 | 0.0039 | 0.00288 | 0.0034 | 0.00576 | 0.02587 | 0.00865 | 0.00349 | 0.01164 | 0.00742 | 0.00461 | 0.01141 | 0.4041 | 0.12131 | 1.25894 | 19.7392 |
| 2-Pyrrolidone-5-carboxylic acid/ L-Pyroglutamic acid | 0.84339 | 0.59407 | 0.49676 | 0.62828 | 0.96012 | 2.70791 | 1.01769 | 0.67871 | 1.12268 | 1.13347 | 0.70452 | 1.33209 | 0.52889 | 0.12285 | 1.25455 | 9.95677 |
| Xanthurenic acid | 0.00143 | 0.00401 | 0.00246 | 0.00439 | 0.00526 | 0.0048 | 0.00464 | 0.0047 | 0.00549 | N/A | 0.00351 | 0.00491 | 0.71522 | 0.12439 | 1.31063 | 16.8596 |
| hypoxanthine | 0.02701 | 0.00957 | 0.0132 | 0.02206 | 0.02246 | 0.02692 | 0.03435 | 0.01965 | 0.02683 | 0.02164 | 0.01886 | 0.02588 | 0.72878 | 0.12637 | 1.24461 | 13.6545 |
| guanine | 0.02163 | 0.03644 | 0.02892 | 0.062 | 0.08442 | 0.05679 | 0.06188 | 0.08437 | 0.06043 | 0.0794 | 0.04668 | 0.06857 | 0.68076 | 0.12793 | 1.24029 | 27.6738 |
| Pyridoxine | 0.02191 | 0.01684 | 0.01987 | 0.02517 | 0.05553 | 0.20066 | 0.0434 | 0.02681 | 0.08356 | 0.05482 | 0.02787 | 0.08185 | 0.34043 | 0.12901 | 1.23728 | 9.68676 |
| Asparagine | 0.01876 | 0.00997 | 0.00937 | 0.01817 | 0.01589 | 0.02319 | 0.01726 | 0.01583 | 0.02423 | 0.05629 | 0.01443 | 0.02736 | 0.52748 | 0.13074 | 1.23253 | 4.34144 |
| 3-Hydroxybenzaldehyde | 0.00601 | 0.00801 | 0.0105 | 0.00847 | 0.007 | 0.01164 | 0.00804 | 0.00801 | 0.00958 | 0.01358 | 0.008 | 0.01017 | 0.78664 | 0.13814 | 1.21263 | 12.4607 |
| taurolithocholate | 0.00363 | 0.00302 | 0.00312 | 0.0065 | 0.00377 | 0.00652 | 0.00416 | 0.00497 | 0.00546 | 0.00504 | 0.00401 | 0.00523 | 0.76655 | 0.14049 | 1.20644 | 9.1598 |
| Uridine 5′-diphospho-N-acetylglucosamine | 0.43163 | 0.14663 | 0.19811 | 0.2407 | 0.28123 | 0.28661 | 0.35654 | 0.27535 | 0.42241 | 0.4408 | 0.25966 | 0.35634 | 0.72868 | 0.14056 | 1.20627 | 11.3294 |
| Arginine | 0.6029 | 0.1588 | 0.23598 | 0.23863 | 0.86235 | 2.23655 | 0.49187 | 0.31085 | 0.94279 | 1.07766 | 0.41973 | 1.01194 | 0.41478 | 0.14141 | 1.20405 | 7.0616 |
| SuccCOA | 0.00013 | 6.4E-05 | 4E-05 | 0.00024 | 0.00023 | 0.00026 | 0.00011 | 0.00027 | 0.00018 | 0.00039 | 0.00014 | 0.00024 | 0.5814 | 0.14173 | 1.20321 | 15.8643 |
| β-Nicotinamide adenine dinucleotide, reduced disodium salt hydrate | 0.25314 | 0.10822 | 0.1102 | 0.12697 | 0.09914 | 0.07303 | 0.11088 | 0.09663 | 0.08227 | 0.09476 | 0.13953 | 0.09151 | 1.52473 | 0.14192 | 1.2027 | 7.92367 |
| S-(5'-Adenosyl)-homocysteine | 0.00515 | 0.00367 | 0.00579 | 0.00865 | 0.00716 | 0.00628 | 0.00755 | 0.01229 | 0.00654 | 0.00905 | 0.00608 | 0.00835 | 0.72906 | 0.14289 | 1.20021 | 10.2701 |
| 2-Hydroxypyridine | 0.00356 | 0.00369 | 0.00178 | 0.00357 | 0.00358 | 0.00259 | 0.00339 | 0.00237 | 0.00211 | 0.00224 | 0.00324 | 0.00254 | 1.27378 | 0.14419 | 1.19683 | 29.5613 |
| Ophthalmate | 0.00324 | 0.00269 | 0.00317 | 0.00474 | 0.00546 | 0.00411 | 0.00425 | 0.00572 | 0.00442 | 0.00659 | 0.00386 | 0.00502 | 0.76966 | 0.1456 | 1.19322 | 22.148 |
| 4-Hydroxy-3-methoxyphenylglycol/ 3,4-Dihydroxymandelic acid | 0.00818 | 0.01149 | 0.01542 | 0.01393 | 0.01271 | 0.01468 | 0.01309 | 0.01618 | 0.01372 | 0.0147 | 0.01235 | 0.01447 | 0.85308 | 0.15003 | 1.18198 | 26.2665 |
| L-Alpha-aminobutyric acid/ 2-Aminoisobutyric acid | 0.01807 | 0.01547 | 0.01709 | 0.03055 | 0.0199 | 0.04509 | 0.0216 | 0.02012 | 0.02634 | 0.02884 | 0.02022 | 0.0284 | 0.71199 | 0.1547 | 1.17032 | 12.613 |
| Tetradecanedioic acid | 0.78828 | 0.92286 | 1.16879 | 0.98851 | 0.86311 | 1.03116 | 0.96963 | 1.16544 | 0.99432 | 1.17782 | 0.94631 | 1.06767 | 0.88633 | 0.15842 | 1.16121 | 11.952 |
| Ornithine | 0.04714 | 0.01914 | 0.03361 | 0.03039 | 0.04159 | 0.07451 | 0.04355 | 0.02911 | 0.05604 | 0.03965 | 0.03437 | 0.04857 | 0.70767 | 0.15947 | 1.15867 | 9.66404 |

| | | | | | | | | | | | | | | | | |
|---|---|---|---|---|---|---|---|---|---|---|---|---|---|---|---|---|
| L-Glutathione reduced | 0.00345 | 0.00337 | 0.00193 | 0.00936 | 0.00772 | 0.00418 | 0.00833 | 0.00815 | 0.00857 | 0.00987 | 0.00517 | 0.00782 | 0.66065 | 0.16108 | 1.15477 | 9.76793 |
| 2'-Deoxyuridine 5'-monophosphate | 0.00681 | N/A | N/A | N/A | 0.00044 | 0.00041 | 0.00038 | 0.00012 | N/A | 0.00025 | 0.00363 | 0.00029 | 12.5476 | 0.16254 | 1.42038 | 21.3226 |
| Adenosine-5'-diphosphoglucose | 0.43642 | 0.37468 | 0.30325 | 0.93195 | 0.91685 | 0.70643 | 0.49968 | 1.45471 | 0.82807 | 1.13119 | 0.59263 | 0.92402 | 0.64136 | 0.16431 | 1.14703 | 17.406 |
| Tyrosine | 0.88215 | 0.2707 | 0.20319 | 0.34117 | 0.39092 | 1.25261 | 0.5757 | 0.29498 | 0.80778 | 0.6457 | 0.41762 | 0.71535 | 0.5838 | 0.17206 | 1.12883 | 2.35413 |
| CDP-ethanolamine | | 1.27168 | 1.00707 | 0.82363 | 1.60914 | 1.5073 | 4.44093 | 1.87862 | 2.022 | 1.16232 | 1.17788 | 2.20224 | 0.53486 | 0.17226 | 1.18567 | 20.8866 |
| Cadaverine | 0.00731 | 0.00648 | 0.008 | 0.00818 | 0.00983 | 0.00882 | 0.00897 | 0.00782 | 0.01147 | 0.00882 | 0.00796 | 0.00918 | 0.86712 | 0.17662 | 1.11834 | 7.82384 |
| 4-Imidazoleacetic acid | 0.0052 | 0.00763 | 0.0041 | 0.00331 | 0.00514 | 0.02931 | 0.00414 | 0.00614 | 0.00835 | 0.01132 | 0.00507 | 0.01185 | 0.42808 | 0.17732 | 1.11675 | 27.7325 |
| OleCOA | 7.6E-05 | 0.00015 | 5.6E-05 | 0.00031 | 0.00014 | 5.3E-05 | 0.00022 | 0.0003 | 0.00041 | 0.0003 | 0.00015 | 0.00026 | 0.574 | 0.18217 | 1.10582 | 14.3481 |
| N α -Acetyl-lysine | 0.03675 | 0.016 | 0.01527 | 0.02614 | 0.02241 | 0.09409 | 0.01809 | 0.02626 | 0.02982 | 0.05061 | 0.02332 | 0.04377 | 0.53265 | 0.18832 | 1.09218 | 7.0848 |
| Hippuric acid | 0.00127 | 0.00113 | 0.0004 | 0.00048 | 0.00087 | 0.00381 | 0.00112 | 0.00077 | 0.0012 | 0.00137 | 0.00083 | 0.00165 | 0.5016 | 0.18892 | 1.09086 | 11.497 |
| 2-Aminocaprylic acid | 0.0088 | 0.00567 | 0.00677 | 0.00712 | 0.00949 | 0.01553 | 0.01105 | 0.00792 | 0.00549 | 0.01097 | 0.00757 | 0.01019 | 0.74298 | 0.18955 | 1.08947 | 9.95681 |
| creatinine | 0.07407 | 0.03268 | 0.02603 | 0.03928 | 0.05316 | 0.14783 | 0.05149 | 0.0356 | 0.06925 | 0.07089 | 0.04505 | 0.07501 | 0.60051 | 0.19358 | 1.08071 | 2.92414 |
| ArachCOA | 4E-05 | 6.9E-05 | 2.6E-05 | 7.4E-05 | 2.3E-05 | 4.3E-05 | 5.2E-05 | 3.6E-05 | 0.00017 | 0.00015 | 4.6E-05 | 8.9E-05 | 0.52048 | 0.19519 | 1.07724 | 28.9226 |
| 7,8-Dihydro-L-biopterin | 0.00072 | 0.00032 | 0.001 | 0.00101 | 0.00127 | 0.00077 | 0.00121 | 0.00087 | 0.00162 | 0.00147 | 0.00086 | 0.00119 | 0.72753 | 0.19908 | 1.06892 | 17.6325 |
| Methylcysteine | 0.00523 | 0.00433 | 0.00237 | 0.00428 | 0.0058 | 0.00718 | 0.00399 | 0.00662 | 0.00473 | 0.00528 | 0.0044 | 0.00556 | 0.7921 | 0.2009 | 1.06506 | 15.3982 |
| Trigonelline | 0.03201 | 0.02762 | 0.03104 | 0.02892 | 0.03073 | 0.04074 | 0.03227 | 0.02689 | 0.03141 | 0.03616 | 0.03007 | 0.03349 | 0.89767 | 0.20192 | 1.06293 | 9.90116 |
| Cytidine 5'-diphosphate sodium salt hydrate | 4.5E-06 | 1.3E-06 | 2.3E-06 | 6.1E-06 | 3E-06 | 3.9E-06 | 9.2E-07 | 5.8E-06 | 1.1E-05 | 1E-05 | 3.4E-06 | 6.3E-06 | 0.55123 | 0.20293 | 1.06079 | 22.2219 |
| Asymmetric dimethylarginine | 0.03799 | 0.0169 | 0.02113 | 0.02509 | 0.02953 | 0.03671 | 0.03052 | 0.02642 | 0.02721 | 0.04298 | 0.02613 | 0.03277 | 0.7974 | 0.20355 | 1.05948 | 4.50466 |
| Succinic acid/ 3-Hydroxy-2-methylbutanoic acid | 0.08177 | 0.07496 | 0.10024 | 0.04363 | 0.06932 | 0.29634 | 0.10779 | 0.05444 | 0.10938 | 0.09737 | 0.07398 | 0.13306 | 0.55599 | 0.20687 | 1.05256 | 7.05297 |
| 3-(2-Hydroxyphenyl)propionic acid | 0.00788 | 0.00231 | 0.00632 | 0.00522 | 0.00275 | 0.00322 | 0.00292 | 0.00499 | 0.00216 | 0.00331 | 0.0049 | 0.00332 | 1.47569 | 0.20859 | 1.04899 | 27.5218 |
| Uridine 5'-monophosphate | 0.06969 | 0.04913 | 0.0034 | N/A | 0.02241 | 0.00735 | 0.08275 | 0.08602 | N/A | 0.14833 | 0.03616 | 0.08111 | 0.44577 | 0.21374 | 1.17298 | 19.3082 |
| Trimethylamine N-oxide | 0.00448 | 0.00324 | 0.00444 | 0.00434 | 0.00322 | 0.00868 | 0.00449 | 0.0038 | 0.00513 | 0.00398 | 0.00394 | 0.00522 | 0.75645 | 0.21454 | 1.03677 | 18.3139 |
| 3,4-Dihydroxymandelic acid | 0.00271 | 0.00365 | 0.00445 | 0.00375 | 0.00361 | 0.00433 | 0.00307 | 0.00445 | 0.00424 | 0.0047 | 0.00363 | 0.00416 | 0.87328 | 0.22067 | 1.0244 | 21.6619 |
| 4-Trimethylammoniobutanoic acid | 0.13299 | 0.09048 | 0.09128 | 0.14201 | 0.14974 | 0.14054 | 0.12118 | 0.12648 | 0.14746 | 0.17621 | 0.1213 | 0.14237 | 0.85198 | 0.2235 | 1.01876 | 8.03391 |
| Indoxyl sulfate | 0.00149 | 0.00316 | 0.0021 | 0.00188 | 0.00219 | 0.00346 | 0.00197 | 0.00228 | 0.00286 | 0.00269 | 0.00216 | 0.00265 | 0.8161 | 0.23069 | 1.00459 | 11.6002 |
| L-Pyroglutamic acid/ 2-Pyrrolidone-5-carboxylic acid | N/A | 0.02105 | 0.02412 | 0.03015 | 0.01819 | 0.01124 | 0.02221 | 0.02577 | 0.00725 | 0.02143 | 0.02338 | 0.01758 | 1.3299 | 0.24766 | 1.02402 | 3.2469 |
| 2,3-Diphosphoglyceric acid | 0.00048 | 0.00077 | 0.0002 | 0.00031 | 0.00048 | 0.00027 | 0.00014 | N/A | N/A | 0.00041 | 0.00045 | 0.00027 | 1.63763 | 0.25923 | 1.04298 | 28.753 |
| Alanine/Sarcosine | 0.09788 | 0.03707 | 0.02796 | 0.04252 | 0.03599 | 0.08396 | 0.06143 | 0.04025 | 0.07396 | 0.07651 | 0.04828 | 0.06722 | 0.71823 | 0.23522 | 0.99579 | 7.5987 |
| 5-(1,2-dithiolan-3-yl)valeramide | 0.00132 | 0.00082 | 0.00096 | 0.00062 | 0.00046 | 0.00062 | 0.00078 | 0.00066 | 0.00055 | 0.00059 | 0.00084 | 0.00064 | 1.30681 | 0.23634 | 0.99361 | 17.0957 |
| Indole-3-carboxylic acid | 0.00026 | 0.00011 | 0.00019 | 0.00022 | 9.2E-05 | 0.00026 | 0.00026 | 0.00029 | 0.00019 | 0.00014 | 0.00017 | 0.00023 | 0.76422 | 0.23831 | 0.98983 | 23.1004 |
| Spermidine | 0.00621 | 0.00381 | 0.00621 | 0.00366 | 0.00484 | 0.00482 | 0.00427 | 0.00413 | 0.00365 | 0.00421 | 0.00495 | 0.00422 | 1.17344 | 0.24664 | 0.97401 | 17.5443 |
| glycine | 0.01062 | 0.00454 | 0.00531 | 0.0069 | 0.00653 | 0.00844 | 0.0076 | 0.00662 | 0.00812 | 0.01118 | 0.00678 | 0.00839 | 0.8079 | 0.24872 | 0.97011 | 10.4564 |
| deoxyinosine | 0.00045 | 4.1E-05 | 2.4E-05 | 4.2E-05 | 5.8E-05 | 3.2E-05 | 2.4E-05 | 6.3E-06 | 2.7E-05 | 1.6E-05 | 0.00012 | 2.1E-05 | 5.78971 | 0.24875 | 0.97006 | 29.6675 |
| Proline | 3.40289 | 1.55354 | 1.66143 | 2.49756 | 2.37515 | 4.20375 | 2.57937 | 1.90886 | 3.00581 | 2.88534 | 2.29811 | 2.91663 | 0.78794 | 0.25207 | 0.96387 | 1.63633 |
| Glutathione oxidized | 0.05536 | 0.06131 | 0.0381 | 0.09529 | 0.09537 | 0.0617 | 0.03652 | 0.06805 | 0.04849 | 0.05294 | 0.06909 | 0.05354 | 1.29043 | 0.25246 | 0.96314 | 5.32681 |
| Alpha-ketoisovaleric acid | 1E-05 | 8.2E-06 | 3.4E-05 | 2.2E-05 | 1.7E-05 | 1.9E-05 | 4.3E-05 | 3.5E-05 | 1E-05 | 3E-05 | 1.8E-05 | 2.7E-05 | 0.66836 | 0.25294 | 0.96224 | 20.4126 |
| Geranyl pyrophosphate | 0.2125 | 0.10002 | 0.23453 | 0.15881 | 0.12133 | 0.12246 | 0.12742 | 0.13776 | 0.11998 | 0.15645 | 0.16544 | 0.13282 | 1.24562 | 0.25475 | 0.95889 | 15.9004 |
| Thymine | 0.03336 | 0.00165 | N/A | 0.00098 | 0.00047 | 0.0013 | 0.00061 | 0.00049 | 0.00061 | 0.00133 | 0.00912 | 0.00087 | 10.5173 | 0.28354 | 0.95661 | 20.9637 |
| Aniline-2-sulfonic acid | 0.00929 | 0.0011 | 0.00108 | 0.00331 | 0.00044 | 0.00012 | 0.00114 | 0.00238 | 0.00077 | 0.00056 | 0.00305 | 0.00099 | 3.06568 | 0.25662 | 0.95544 | 18.4516 |
| Ethyl 3-indoleacetate | 0.00017 | 0.00011 | 0.00018 | 0.00044 | 0.00023 | 0.0002 | 2.1E-05 | 0.00013 | 0.00014 | 0.00023 | 0.00023 | 0.00014 | 1.56056 | 0.25954 | 0.95009 | 23.1049 |
| Nicotinic acid adenine dinucleotide | 0.01096 | 0.00311 | 0.00285 | 0.00548 | 0.00623 | 0.00773 | 0.00569 | 0.00698 | 0.00931 | 0.00843 | 0.00573 | 0.00763 | 0.75076 | 0.26582 | 0.93868 | 17.2268 |
| Palmitoleic acid | 0.00021 | 7.1E-06 | 6.9E-05 | 0.00026 | 0.00015 | 7.7E-05 | 8.1E-05 | 5.7E-05 | 4.3E-06 | 0.00016 | 0.00014 | 7.7E-05 | 1.824 | 0.26596 | 0.93842 | 22.1033 |
| Acetyl carnitine | 0.22476 | 0.10926 | 0.09217 | 0.19794 | 0.16846 | 0.13491 | 0.15075 | 0.21492 | 0.22591 | 0.28881 | 0.15852 | 0.20306 | 0.78066 | 0.27013 | 0.93094 | 4.84734 |
| Homoarginine | 0.00545 | 0.00377 | 0.00301 | 0.0025 | 0.00303 | 0.00928 | 0.00375 | 0.00142 | 0.007 | 0.00479 | 0.00355 | 0.00525 | 0.67619 | 0.27314 | 0.92556 | 18.2129 |
| Nicotinamide adenine dinucleotide phosphate | 3.9E-05 | 0.00011 | 2.3E-05 | 0.0003 | 0.00016 | 0.00022 | 0.00013 | 0.00051 | 1.6E-05 | 0.00033 | 0.00013 | 0.00024 | 0.52037 | 0.27433 | 0.92346 | 27.3699 |
| 3-Methoxy-tyrosine | 0.24538 | 0.12342 | 0.19531 | 0.27053 | 0.1728 | 0.27026 | 0.63831 | 0.1836 | 0.21456 | 0.21923 | 0.20149 | 0.30519 | 0.6602 | 0.27438 | 0.92336 | 6.57603 |
| Pyridoxamine | 0.00736 | 0.00314 | 0.00287 | 0.00418 | 0.0058 | 0.01669 | 0.00611 | 0.00298 | 0.00678 | 0.00546 | 0.00467 | 0.0076 | 0.61444 | 0.27587 | 0.92072 | 20.389 |
| 4-Hydroxybenzoic acid | 0.00033 | 8.4E-05 | 0.00059 | 0.00043 | 0.00031 | 0.00022 | 0.00047 | 0.00053 | 0.00043 | 0.00084 | 0.00035 | 0.0005 | 0.70267 | 0.2853 | 0.90421 | 23.9138 |
| Histidine | 0.07715 | 0.01243 | 0.00887 | 0.01059 | 0.02172 | 0.11838 | 0.02855 | 0.01551 | 0.06814 | 0.02982 | 0.02615 | 0.05208 | 0.50214 | 0.28821 | 0.89916 | 24.5901 |
| Hexose-phosphate | 0.41983 | 0.12659 | 0.14395 | 0.26276 | 0.1526 | 0.07501 | 0.16465 | 0.1824 | 0.11178 | 0.2283 | 0.22115 | 0.15243 | 1.45085 | 0.29509 | 0.88735 | 11.7456 |
| Oxalomalate | 0.00943 | 0.00213 | 0.00423 | 0.00249 | 0.00518 | 0.01131 | 0.00629 | 0.00429 | 0.00571 | 0.00583 | 0.00469 | 0.00668 | 0.70207 | 0.2954 | 0.88682 | 29.6086 |
| Succinic acid semialdehyde | 0.01217 | 0.00753 | 0.01074 | 0.00825 | 0.00811 | 0.00983 | 0.01175 | 0.00847 | 0.01218 | 0.01068 | 0.00936 | 0.01058 | 0.88457 | 0.30543 | 0.86988 | 18.4601 |
| Linoleic acid | 0.0007 | 0.00121 | 0.00055 | 0.00012 | 0.00024 | 0.00188 | 0.00143 | 9.7E-05 | 0.00128 | 0.00028 | 0.00056 | 0.00099 | 0.56702 | 0.30698 | 0.86728 | 16.2784 |
| Threonine/Homoserine | 0.09793 | 0.03377 | 0.03164 | 0.041 | 0.04562 | 0.09919 | 0.0611 | 0.03296 | 0.08203 | 0.06391 | 0.04999 | 0.06784 | 0.73695 | 0.31161 | 0.85959 | 9.29348 |
| Cholesteryl acetate | 0.32345 | 0.39694 | 0.44209 | 0.45299 | 0.40411 | 0.42309 | 0.39287 | 0.47308 | 0.43151 | 0.43855 | 0.40391 | 0.43182 | 0.93537 | 0.31791 | 0.84919 | 14.0685 |
| Hydrocortisone 21-acetate | 0.01977 | 0.01555 | 0.00186 | 0.01379 | 0.00484 | 0.00276 | 0.01735 | 0.00631 | 0.00475 | 0.00118 | 0.01116 | 0.00647 | 1.72483 | 0.31905 | 0.84733 | 19.8889 |
| Stearic acid | 0.00349 | 0.00138 | 0.00103 | 0.00119 | 0.00123 | 0.00089 | 0.00139 | 0.00106 | 0.00111 | 0.00138 | 0.00166 | 0.00117 | 1.4287 | 0.31954 | 0.84652 | 16.1317 |
| Xanthosine | 0.02872 | 0.00285 | 0.00265 | 0.00455 | 0.00185 | 0.00313 | 0.00242 | 0.00167 | 0.00322 | 0.00282 | 0.00812 | 0.00265 | 3.06281 | 0.32136 | 0.84356 | 15.9646 |
| L-Cysteic acid | 0.00601 | 0.00193 | 0.00297 | 0.00165 | 0.00092 | 0.0024 | 0.00123 | 0.00128 | 0.00137 | 0.00231 | 0.0027 | 0.00172 | 1.57037 | 0.32241 | 0.84184 | 20.1148 |
| Phosphocholine | 0.27896 | 0.23376 | 0.31724 | 0.46008 | 0.42085 | 0.2887 | 0.39434 | 0.41036 | 0.44267 | 0.44188 | 0.34218 | 0.39559 | 0.86499 | 0.32754 | 0.83353 | 7.00375 |
| Phosphocreatine | 0.02151 | 0.00821 | 0.00432 | 0.0273 | 0.00752 | 0.01389 | 0.01607 | 0.01427 | 0.02339 | 0.02935 | 0.01377 | 0.01939 | 0.71026 | 0.32921 | 0.83083 | 10.2296 |
| N1-Acetylspermine | 0.00241 | 0.00182 | 0.0032 | 0.00222 | 0.00275 | 0.00239 | 0.00788 | 0.00165 | 0.00365 | 0.00267 | 0.00248 | 0.00365 | 0.67928 | 0.33017 | 0.82929 | 23.1214 |
| beta aminobutyric acid | 0.00397 | 0.00386 | 0.00386 | 0.00708 | 0.0039 | 0.00791 | 0.003 | 0.00515 | 0.00682 | 0.00522 | 0.00454 | 0.00562 | 0.80711 | 0.33199 | 0.82636 | 21.7315 |
| Tryptophanol | 0.00069 | 0.00099 | 0.00123 | 0.00112 | 0.00126 | 0.00142 | 0.00111 | 0.00163 | 0.00116 | 0.00084 | 0.00106 | 0.00123 | 0.85847 | 0.33236 | 0.82578 | 24.2843 |
| dimethylglycine | 0.12497 | 0.08903 | 0.11477 | 0.18502 | 0.15293 | 0.2154 | 0.1403 | 0.10732 | 0.17177 | 0.15519 | 0.13334 | 0.158 | 0.84396 | 0.33995 | 0.81369 | 11.53 |
| Methylmalonic acid | 0.00246 | 0.00152 | 0.0027 | 0.00211 | 0.00236 | 0.00299 | 0.00195 | 0.00342 | 0.002 | 0.00255 | 0.00223 | 0.00258 | 0.8637 | 0.34108 | 0.8119 | 17.8924 |
| Palmitoylcarnitine (Carnitine_C16) | N/A | 0.01895 | 0.01782 | 0.03366 | 0.02149 | 0.02317 | 0.04351 | 0.0178 | 0.03419 | 0.02439 | 0.02298 | 0.02861 | 0.8032 | 0.38587 | 0.78524 | 10.2143 |
| Serine | 0.05684 | 0.04451 | 0.02694 | 0.0254 | 0.0224 | 0.05388 | 0.0421 | 0.02695 | 0.05183 | 0.04114 | 0.03522 | 0.04318 | 0.81558 | 0.35933 | 0.78342 | 16.2551 |
| Cysteinesulfinic acid | 0.01195 | 0.00113 | 0.00107 | 0.00543 | 0.00125 | 0.00146 | 0.00094 | 0.00517 | 0.00152 | 0.0008 | 0.00416 | 0.00198 | 2.10607 | 0.36261 | 0.77838 | 11.3064 |
| Palmitoyl-DL-carnitine chloride | 0.01825 | 0.00042 | 0.00036 | 0.0007 | 0.00054 | 0.00032 | 0.00124 | 0.00037 | 0.00084 | 0.0004 | 0.00405 | 0.00064 | 6.36648 | 0.36452 | 0.77545 | 7.17061 |
| mono-Ethyl malonate | 0.01428 | 0.0086 | 0.00579 | 0.00949 | 0.00611 | 0.0135 | 0.01098 | 0.00491 | 0.01755 | 0.00977 | 0.00885 | 0.01134 | 0.78071 | 0.36489 | 0.7749 | 22.2504 |
| Stearoyl-L-carnitine | 0.02339 | 7.1E-05 | 9.4E-05 | 9.2E-05 | 7.4E-05 | 7.8E-06 | 0.00102 | 6.5E-05 | 0.00014 | 0.00011 | 0.00474 | 0.00027 | 17.6687 | 0.36548 | 0.77399 | 11.1827 |
| Anthranilic acid | 0.00442 | 0.00145 | 0.00345 | 0.00271 | 0.00173 | 0.00242 | 0.00265 | 0.00239 | 0.00128 | 0.00216 | 0.00275 | 0.00218 | 1.26215 | 0.3668 | 0.77197 | 28.6113 |
| 4-Acetamidobutanoic acid | 0.01052 | 0.01119 | 0.01051 | 0.01109 | 0.01183 | 0.01122 | 0.00873 | 0.01163 | 0.01318 | 0.01692 | 0.01103 | 0.01234 | 0.89395 | 0.3683 | 0.76969 | 11.3653 |
| Xanthosine 5'-monophosphate (sodium salt) | 3.6E-06 | N/A | 5.6E-06 | 3.3E-06 | 4.6E-06 | 3.7E-06 | N/A | N/A | 3.7E-06 | N/A | 4.3E-06 | 3.7E-06 | 1.15393 | 0.49812 | 0.76233 | 29.5356 |
| Myristoyl-L-carnitine | 0.00973 | 0.00021 | 0.00012 | 0.00027 | 0.00016 | 9.5E-05 | 0.00084 | 0.00017 | 0.00031 | 0.00018 | 0.0021 | 0.00032 | 6.58941 | 0.37926 | 0.75314 | 3.26166 |
| Tricosanoic acid | 0.40852 | 0.49567 | 0.7709 | 0.58902 | 0.59244 | 0.60706 | 0.54022 | 0.64813 | 0.60622 | 0.80094 | 0.57131 | 0.64051 | 0.89196 | 0.38004 | 0.75197 | 6.50456 |
| 2-Aminoethylphosphonic acid | 0.00408 | 0.00122 | 0.00193 | 0.00221 | 0.00183 | 0.00108 | 0.00196 | 0.00154 | 0.00213 | 0.00213 | 0.00226 | 0.00177 | 1.27554 | 0.38092 | 0.75066 | 16.2622 |
| 5-Aminovaleric acid/ Guanidineacetic acid | 0.03595 | 0.01859 | 0.01914 | 0.02751 | 0.01348 | 0.01904 | 0.01891 | 0.01809 | 0.0203 | 0.02103 | 0.02293 | 0.01948 | 1.1775 | 0.41162 | 0.70558 | 11.6322 |
| 2'-Deoxyadenosine 5'-triphosphate | 0.86996 | 0.82317 | 0.7254 | 0.93464 | 0.88068 | 0.93376 | 0.83685 | 1.01352 | 0.91601 | 0.76897 | 0.84677 | 0.89382 | 0.94736 | 0.41513 | 0.70052 | 7.542 |
| Phenylpyruvate | 0.00115 | 0.00062 | 0.00052 | 0.0004 | 0.00053 | 0.00102 | 0.00056 | 0.00025 | 0.00039 | 0.00013 | 0.00064 | 0.00047 | 1.36686 | 0.42084 | 0.69236 | 18.089 |
| O-Phospho-serine | 0.00673 | 0.00445 | 0.00511 | 0.00836 | 0.0092 | 0.00535 | 0.00711 | 0.00779 | 0.00834 | 0.01044 | 0.00677 | 0.0078 | 0.86772 | 0.42618 | 0.68477 | 14.6705 |
| Phenethylamine | 0.00272 | 0.00269 | 0.00321 | 0.00192 | 0.00245 | 0.00195 | 0.00277 | 0.00227 | 0.00254 | 0.00244 | 0.0026 | 0.00239 | 1.08593 | 0.43453 | 0.67297 | 17.7475 |
| Arabitol | 0.00031 | 0.00013 | 0.00026 | 0.00022 | 0.00016 | 0.00023 | 0.00019 | 0.00016 | 0.00011 | 0.00022 | 0.00022 | 0.00018 | 1.17829 | 0.43675 | 0.66986 | 20.7469 |
| Citrulline | 0.17472 | 0.0318 | 0.00548 | 0.01575 | 0.00685 | 0.03414 | 0.02619 | 0.00914 | 0.01026 | 0.0219 | 0.04692 | 0.02033 | 2.3083 | 0.43885 | 0.66692 | 10.0308 |
| Glycerol 3-phosphate | 0.1162 | 0.06705 | 0.1026 | 0.18813 | 0.13734 | 0.10863 | 0.15031 | 0.09514 | 0.21225 | 0.16193 | 0.12226 | 0.14565 | 0.83943 | 0.44146 | 0.66327 | 28.9616 |
| Lauroyl-L-carnitine | 0.00212 | 5.8E-05 | 4.5E-05 | 0.00013 | 2.7E-05 | 9.8E-05 | 0.00019 | 5.5E-05 | 0.00026 | 0.00011 | 0.00048 | 0.00014 | 3.34222 | 0.4435 | 0.66042 | 10.5467 |
| Decanoyl-L-carnitine | 0.00085 | 2.5E-05 | 4.3E-05 | 5.2E-05 | 4.7E-05 | 1.5E-05 | 8.9E-05 | 2.2E-05 | 0.00015 | 8.1E-05 | 0.0002 | 7.2E-05 | 2.84111 | 0.44428 | 0.65934 | 24.5303 |
| Adenosine triphosphate | 0.52915 | 0.3991 | 0.06835 | 1.61961 | N/A | 0.5133 | 0.39062 | 1.37598 | N/A | 1.61001 | 0.65405 | 0.97248 | 0.67256 | 0.50944 | 0.65388 | 22.2823 |
| 1-Hydroxy-2-naphthoic acid | 6.1E-05 | 0.00059 | 0.00021 | 2.3E-05 | 0.00052 | 0.00063 | 0.00049 | 0.00057 | 0.00032 | 3.4E-05 | 0.00028 | 0.00041 | 0.69085 | 0.44882 | 0.65303 | 7.03058 |
| acetyl phosphate | 0.03522 | 0.045 | 0.04295 | 0.05335 | 0.04561 | 0.04458 | 0.04691 | 0.04434 | 0.05102 | 0.0477 | 0.04443 | 0.04691 | 0.94707 | 0.45265 | 0.64774 | 10.9968 |

| | | | | | | | | | | | | | | | | |
|---|---|---|---|---|---|---|---|---|---|---|---|---|---|---|---|---|
| Adenosine | 0.00955 | 0.00417 | 0.00299 | 0.00499 | 0.00572 | 0.00426 | 0.00455 | 0.00869 | 0.00728 | 0.00838 | 0.00548 | 0.00663 | 0.82689 | 0.4532 | 0.64699 | 9.18046 |
| Trisodium 2-metylcitrate, racemic mixture | 0.00026 | 0.00034 | 0.00029 | 0.00024 | 0.00024 | 0.00024 | 0.00011 | 0.00012 | 0.00042 | N/A | 0.00027 | 0.00022 | 1.22162 | 0.47923 | 0.6469 | 28.1849 |
| Homogentisic acid | 0.00089 | 0.00049 | 0.00133 | 0.00112 | 0.00096 | N/A | 0.00107 | 0.00097 | N/A | 0.00126 | 0.00096 | 0.0011 | 0.87256 | 0.50336 | 0.64187 | 18.0446 |
| 3-Dehydroshikimic acid | 1.57461 | 0.20816 | 0.32601 | 0.19396 | 0.19277 | 0.43447 | 0.20718 | 0.23006 | 0.23982 | 0.31715 | 0.4991 | 0.28574 | 1.74671 | 0.4573 | 0.64135 | 27.1557 |
| Erythrose 4-phosphate | 0.0131 | 0.00514 | 0.00834 | 0.00723 | 0.00508 | 0.00446 | 0.00914 | 0.00554 | 0.0056 | 0.00755 | 0.00778 | 0.00646 | 1.20432 | 0.45751 | 0.64106 | 17.3007 |
| Quinaldic acid | 0.00134 | 0.00063 | 0.00071 | 0.00126 | 0.00047 | 0.00037 | 0.00056 | 0.0006 | 0.00113 | 0.00089 | 0.00088 | 0.00071 | 1.24283 | 0.46037 | 0.63714 | 25.5355 |
| (S)-Mevalonic acid | 0.04723 | 0.00655 | 0.01816 | 0.00941 | 0.00369 | 0.01017 | N/A | 0.00836 | 0.00548 | 0.0169 | 0.01701 | 0.01023 | 1.66309 | 0.48712 | 0.63577 | 18.8919 |
| Phosphocholine | 6.38787 | 3.85097 | 6.74605 | 8.46581 | 7.99633 | 5.44799 | 6.87053 | 7.9379 | 8.06081 | 9.03035 | 6.68941 | 7.46951 | 0.89556 | 0.46276 | 0.63387 | 6.21161 |
| meso-Tartaric acid | 0.00334 | 0.00174 | 0.00115 | 0.00159 | 0.00076 | 0.00286 | 0.00193 | 0.0015 | 0.0022 | 0.00198 | 0.00172 | 0.0021 | 0.81907 | 0.46535 | 0.63033 | 27.5741 |
| Erucic acid | 6.01572 | 9.28009 | 20.5774 | 13.9171 | 10.0219 | 12.923 | 16.4145 | 13.8749 | 13.1186 | 13.2785 | 11.9624 | 13.9219 | 0.85925 | 0.46855 | 0.62598 | 22.854 |
| Gluconic acid | 0.12836 | 0.0089 | 0.00501 | 0.00577 | 0.00921 | 0.02101 | 0.01125 | 0.00761 | 0.01235 | 0.0127 | 0.03145 | 0.01299 | 2.42188 | 0.4699 | 0.62415 | 17.9148 |
| 1-Aminocyclopropanecarboxylic acid | 0.05043 | 0.0371 | 0.02472 | 0.04851 | 0.03685 | 0.05788 | 0.03401 | 0.02969 | 0.04127 | 0.06697 | 0.03952 | 0.04596 | 0.8598 | 0.47016 | 0.62379 | 23.2014 |
| L-α-Phosphatidylcholine | 0.00035 | 3E-06 | 6.2E-06 | 4E-06 | 4.3E-06 | 7.4E-05 | 1.9E-06 | 1.6E-05 | 9.5E-06 | 1.4E-06 | 7.3E-05 | 2.1E-05 | 3.57135 | 0.47405 | 0.61853 | 25.7766 |
| N-acetyltryptophan | 0.0006 | 0.00041 | 0.00056 | 0.00069 | 0.00045 | 0.00086 | 0.00054 | 0.00041 | 0.00063 | 0.00059 | 0.00054 | 0.00061 | 0.89038 | 0.47476 | 0.61757 | 20.6897 |
| Aminoadipic acid | 0.05087 | 0.00521 | 0.00362 | 0.00953 | 0.00262 | 0.00653 | 0.00065 | 0.00228 | 0.01278 | 0.01377 | 0.01437 | 0.0072 | 1.99484 | 0.47581 | 0.61616 | 27.1204 |
| 8-Hydroxy-2′-deoxyguanosine | 0.00111 | 0.00062 | 0.00084 | 0.00064 | 0.00049 | 0.00042 | 0.00059 | 0.0008 | 0.00071 | 0.00071 | 0.00074 | 0.00065 | 1.14531 | 0.48057 | 0.60975 | 16.436 |
| Fumaric acid | 0.01668 | 0.00161 | N/A | 0.00224 | 0.00154 | 0.00291 | 0.00231 | 0.00278 | 0.00432 | N/A | 0.00552 | 0.00308 | 1.7914 | 0.5397 | 0.60949 | 20.3958 |
| Hypotaurine | 0.0165 | 0.00766 | 0.01341 | 0.01257 | 0.01339 | 0.01273 | 0.01398 | 0.01378 | 0.01277 | 0.01595 | 0.01271 | 0.01384 | 0.91791 | 0.48267 | 0.60692 | 3.75789 |
| L-Methionine sulfoxide | 0.01259 | 0.00038 | 0.00031 | 0.00055 | 0.0003 | 0.00203 | 0.00083 | 0.00029 | 0.0009 | 0.00107 | 0.00282 | 0.00102 | 2.76518 | 0.48414 | 0.60495 | 11.2437 |
| cAMP_pos | 0.0033 | 0.00164 | 0.00252 | 0.00319 | 0.00219 | 0.00131 | 0.00275 | 0.00343 | 0.00126 | 0.00219 | 0.00257 | 0.00219 | 1.1742 | 0.48428 | 0.60477 | 14.3178 |
| Imidazole | 0.00137 | 0.00452 | 0.00319 | 0.00163 | 0.00334 | 0.01648 | 0.00147 | 0.00158 | 0.002 | 0.00322 | 0.00281 | 0.00495 | 0.56754 | 0.48979 | 0.5974 | 26.2367 |
| N-Acetyl-D-Glucosamine 6-Phosphate | 0.05878 | 0.01831 | 0.03223 | 0.04063 | 0.03627 | 0.03763 | 0.04842 | 0.04195 | 0.0347 | 0.04936 | 0.03724 | 0.04241 | 0.87813 | 0.49115 | 0.59559 | 8.63961 |
| Pipecolinic acid | N/A | N/A | N/A | N/A | 0.93996 | 3.88142 | 0.88638 | N/A | 1.50623 | 1.1431 | 0.93996 | 1.85429 | 0.50691 | #DIV/0! | 0.59021 | 10.6773 |
| Stachyose | 0.80194 | 0.2154 | 0.18121 | 0.3672 | 0.32191 | 0.34083 | 0.19416 | 0.23629 | 0.3039 | 0.39606 | 0.37753 | 0.29425 | 1.28305 | 0.49707 | 0.58773 | 27.7025 |
| Homocysteic acid | 0.00712 | 0.00232 | 0.00268 | 0.00161 | 0.00194 | 0.00311 | 0.00258 | 0.00208 | 0.00191 | 0.00234 | 0.00314 | 0.0024 | 1.30415 | 0.49955 | 0.58445 | 26.2539 |
| 3-Phenyllactic acid | 0.01339 | 0.00121 | 0.00183 | 0.00201 | 0.00184 | 0.00296 | 0.00266 | 0.0021 | 0.00228 | 0.00227 | 0.00406 | 0.00245 | 1.65314 | 0.51314 | 0.5666 | 17.0089 |
| Taurine | 9.20109 | 1.51839 | 2.27046 | 3.18317 | 2.46639 | 2.07827 | 2.76261 | 2.83184 | 2.69438 | 3.48928 | 3.7279 | 2.77127 | 1.34519 | 0.5171 | 0.56144 | 8.01225 |
| Mandelic acid | 0.01584 | 0.0071 | 0.0077 | 0.00724 | 0.00666 | 0.00753 | 0.00722 | 0.00752 | 0.00807 | 0.00833 | 0.00891 | 0.00773 | 1.15167 | 0.52203 | 0.55503 | 13.9316 |
| Glutamic acid | 1.49655 | 0.56264 | 0.81248 | 2.09917 | 1.17909 | 1.79622 | 1.08772 | 1.00729 | 1.14323 | 2.36495 | 1.22999 | 1.47988 | 0.83114 | 0.52464 | 0.55165 | 5.68512 |
| Citramalate | 0.13792 | 0.14091 | 0.14349 | 0.16688 | 0.13743 | 0.14531 | 0.14312 | 0.13236 | 0.15121 | 0.13266 | 0.14532 | 0.14093 | 1.03116 | 0.52562 | 0.55038 | 16.3564 |
| FADH | 0.00012 | 0.00013 | 0.00014 | 0.00026 | 0.0003 | 0.0002 | 0.00023 | 1E-04 | 0.00028 | 0.00032 | 0.00019 | 0.00022 | 0.84912 | 0.53129 | 0.54306 | 29.6407 |
| Homoserine | 0.10337 | 0.04129 | 0.0252 | 0.03357 | 0.04872 | 0.09294 | 0.0597 | 0.02882 | 0.06583 | 0.05868 | 0.05043 | 0.06119 | 0.82409 | 0.54815 | 0.52148 | 3.32654 |
| Hexanoyl-L-carnitine (Carnitine_C6) | 0.00092 | 0.0007 | 0.00052 | 0.00056 | 0.00057 | 0.00122 | 0.00012 | 0.00078 | 0.00101 | 0.00076 | 0.00065 | 0.00078 | 0.84147 | 0.5498 | 0.51938 | 13.4157 |
| Serotonin | 0.00038 | 0.00011 | 0.00067 | 0.00027 | 0.00012 | 0.00024 | 0.00042 | 0.00011 | 0.00033 | 9.3E-05 | 0.00031 | 0.00024 | 1.31772 | 0.55136 | 0.5174 | 28.093 |
| 5-Aminolevulinic acid | 0.01531 | 0.00369 | 0.00386 | 0.00537 | 0.00398 | 0.00614 | 0.00349 | 0.00445 | 0.0056 | 0.00542 | 0.00644 | 0.00502 | 1.28309 | 0.55148 | 0.51725 | 19.6019 |
| Riboflavin | 0.00197 | 0.00144 | 0.0018 | 0.00385 | 0.00347 | 0.00329 | 0.00275 | 0.00142 | 0.00486 | 0.0025 | 0.00251 | 0.00296 | 0.8467 | 0.55745 | 0.50969 | 3.26674 |
| Coenzyme A | 3.4E-06 | 2.1E-06 | 9.3E-07 | 1.3E-05 | 2.2E-06 | 3.1E-06 | 2.1E-06 | 5.5E-06 | 1.8E-06 | 2.4E-06 | 4.4E-06 | 3E-06 | 1.47602 | 0.562 | 0.50395 | 29.031 |
| Pentose-phosphate | 0.05165 | 0.01709 | 0.01573 | 0.02441 | 0.01404 | 0.01235 | 0.07347 | 0.01905 | 0.03245 | 0.02363 | 0.02458 | 0.03219 | 0.7637 | 0.57131 | 0.49226 | 9.38131 |
| 5β-Cholanoic acid-3α, 6α, 7β-triol | 0.02679 | 0.03575 | N/A | 0.05122 | 0.02873 | 0.03422 | 0.03592 | 0.03786 | 0.04175 | 0.04277 | 0.03562 | 0.0385 | 0.92522 | 0.59813 | 0.48606 | 15.2402 |
| 2'-Deoxyadenosine 5'-monophosphate | 0.00201 | 0.00084 | 0.00209 | 0.00219 | 0.00254 | 0.00188 | 0.00183 | 0.00217 | 0.00205 | 0.00266 | 0.00193 | 0.00212 | 0.91126 | 0.57769 | 0.48429 | 13.0631 |
| Salicylamide | 0.00045 | 0.00047 | 0.00016 | 0.00037 | 0.00028 | 0.00039 | 0.00017 | 0.00055 | 0.00029 | 0.00061 | 0.00035 | 0.0004 | 0.85801 | 0.58083 | 0.48037 | 17.637 |
| 4-Hydroxybenzaldehyde | 0.00988 | 0.00544 | 0.00502 | 0.00441 | 0.00387 | 0.00559 | 0.00416 | 0.0044 | 0.00346 | 0.0074 | 0.00572 | 0.005 | 1.14427 | 0.58733 | 0.47231 | 18.1791 |
| 4-Imidazoleacrylic acid | 0.00996 | 0.05195 | 0.00267 | 0.01642 | 0.01422 | 0.01048 | 0.02449 | 0.00995 | 0.01123 | 0.01378 | 0.01904 | 0.01398 | 1.36169 | 0.58839 | 0.47099 | 13.9672 |
| 5-Hydroxyindoleacetic acid | 0.02953 | 0.01393 | 0.01269 | 0.01089 | 0.01125 | 0.01315 | 0.01261 | 0.01616 | 0.01199 | 0.01433 | 0.01566 | 0.01365 | 1.14724 | 0.59064 | 0.46821 | 23.495 |
| Petroselinic acid | 0.04998 | 0.06259 | 0.13091 | 0.09164 | 0.06628 | 0.08198 | 0.09266 | 0.10379 | 0.08795 | 0.07565 | 0.08028 | 0.08841 | 0.90805 | 0.6057 | 0.44968 | 19.879 |
| guanosine | 0.02518 | 0.01017 | 0.0133 | 0.01939 | 0.02144 | 0.01362 | 0.02116 | 0.01831 | 0.01896 | 0.02659 | 0.0179 | 0.01973 | 0.90712 | 0.60942 | 0.44513 | 21.4983 |
| Caffeine | 0.00482 | 0.00201 | 0.00255 | 0.00246 | 0.00222 | 0.00273 | 0.00261 | 0.0024 | 0.00202 | 0.00291 | 0.00281 | 0.00253 | 1.1105 | 0.61427 | 0.43921 | 13.7284 |
| β-Nicotinamide adenine dinucleotide | 0.08194 | 0.04448 | 0.09036 | 0.07606 | 0.06139 | 0.0551 | 0.06381 | 0.07129 | 0.06095 | 0.07926 | 0.07085 | 0.06608 | 1.07209 | 0.61614 | 0.43692 | 5.70632 |
| hexol_Dulcitol/Sorbitol | 0.01267 | 0.00325 | 0.00414 | 0.00307 | 0.00498 | 0.01152 | 0.00491 | 0.00348 | 0.00606 | 0.00783 | 0.00562 | 0.00676 | 0.83154 | 0.62919 | 0.42109 | 18.2104 |
| Methylguanidine | 0.0152 | 0.00669 | N/A | 0.00725 | 0.00215 | 0.01179 | 0.00402 | 0.00284 | 0.00403 | 0.00908 | 0.00782 | 0.00635 | 1.23079 | 0.64933 | 0.42045 | 15.7242 |
| Lactic acid | 0.40841 | 0.10596 | 0.12098 | 0.0751 | 0.0767 | 0.17609 | 0.18079 | 0.09169 | 0.0641 | 0.10587 | 0.15743 | 0.12371 | 1.2726 | 0.63088 | 0.41905 | 15.3212 |
| 4-Hydroxy-3-methoxymandelic acid | 0.00298 | 0.00296 | 0.00382 | 0.00472 | 0.00314 | 0.00275 | 0.00286 | 0.00286 | 0.00437 | 0.00365 | 0.00352 | 0.0033 | 1.06881 | 0.63488 | 0.41422 | 27.5914 |
| 3-hydroxybutyric acid | 0.01114 | 0.00102 | 0.0025 | 0.00443 | 0.005 | 0.00346 | 0.00606 | 0.00154 | 0.00599 | 0.00223 | 0.00482 | 0.00386 | 1.24864 | 0.6391 | 0.40914 | 14.8912 |
| 5β-Cholanic acid-3α-ol | 0.56248 | 0.54134 | 0.66898 | 0.58014 | 0.4727 | 0.55635 | 0.51238 | 0.64069 | 0.54404 | 0.67344 | 0.56513 | 0.58538 | 0.96541 | 0.65801 | 0.38651 | 13.5881 |
| Valeryl-carnitine/ Isovaleryl-carnitine (Carnitine_C5) | 0.02502 | 0.01261 | 0.01433 | 0.01585 | 0.01329 | 0.02127 | 0.0157 | 0.01462 | 0.01727 | 0.01807 | 0.01622 | 0.01739 | 0.93289 | 0.65807 | 0.38644 | 12.4747 |
| Flavin adenine dinucleotide | 0.05283 | 0.0257 | 0.04825 | 0.04045 | 0.03554 | 0.03105 | 0.04829 | 0.03964 | 0.04522 | 0.05228 | 0.04055 | 0.0433 | 0.93667 | 0.66166 | 0.38216 | 4.93653 |
| Uracil | 0.01273 | 0.00259 | 0.00386 | 0.00209 | 0.00382 | 0.00887 | 0.00876 | 0.00383 | 0.00468 | 0.00409 | 0.00502 | 0.00605 | 0.83021 | 0.66236 | 0.38133 | 16.4836 |
| Dihydroxyfumaric acid | 0.11416 | 0.11137 | 0.11576 | 0.13036 | 0.10817 | 0.11663 | 0.11103 | 0.11299 | 0.12169 | 0.10725 | 0.11596 | 0.11392 | 1.01796 | 0.6648 | 0.37844 | 16.3115 |
| 1,7-Dimethylxanthine/ Theophylline | 0.0026 | 0.00141 | 0.00174 | 0.00102 | 0.0012 | 0.00175 | 0.0014 | 0.0014 | 0.00077 | 0.00188 | 0.00159 | 0.00144 | 1.10498 | 0.66708 | 0.37573 | 23.0526 |
| Retinoic acid | 0.00066 | 0.00039 | 0.0017 | 0.00076 | 0.00098 | 0.00081 | 0.00228 | 0.00079 | 0.00037 | 0.00108 | 0.0009 | 0.00107 | 0.84218 | 0.67915 | 0.36145 | 28.9975 |
| 6-Phosphogluconic acid | N/A | 0.00712 | 0.00715 | 0.01329 | 0.01302 | 0.01611 | 0.009 | 0.01519 | 0.01242 | 0.01639 | 0.01014 | 0.01382 | 0.73396 | 0.13817 | 0.35657 | 18.3944 |
| 2,6-Diaminoheptandioate | 0.00165 | 0.00071 | 0.00044 | 0.00052 | 0.00043 | 0.00095 | 0.00036 | 0.00069 | 0.00066 | 0.0006 | 0.00075 | 0.00065 | 1.15557 | 0.6953 | 0.34247 | 24.7914 |
| cholate | 0.07815 | 0.06724 | 0.20961 | 0.19726 | 0.05314 | 0.08797 | 0.16041 | 0.08246 | 0.10879 | 0.09133 | 0.12108 | 0.10619 | 1.14021 | 0.69626 | 0.34136 | 15.6477 |
| Guanidine | 0.12627 | 0.06288 | 0.05995 | 0.0596 | 0.06216 | 0.08837 | 0.05118 | 0.06455 | 0.06963 | 0.06925 | 0.07417 | 0.0686 | 1.08127 | 0.70761 | 0.3281 | 9.66631 |
| β-Nicotinamide mononucleotide | 5.7E-06 | 1.2E-05 | 1.9E-05 | 1E-05 | 1.9E-05 | 3.8E-06 | 1.7E-05 | 7.6E-06 | 1.8E-05 | N/A | 1.3E-05 | 1.2E-05 | 1.13374 | 0.72562 | 0.32557 | 17.3559 |
| O-Phospho-L-serine | 0.00629 | 0.0018 | 0.00088 | 0.00478 | 0.00133 | 0.00286 | 0.00356 | 0.00264 | 0.00362 | 0.00449 | 0.00302 | 0.00343 | 0.87836 | 0.71728 | 0.31686 | 11.8202 |
| 3-Methylhistamine/ 1-Methylhistamine | 0.00646 | 0.00341 | 0.00576 | 0.00561 | 0.00473 | 0.00454 | 0.00533 | 0.00477 | 0.00593 | 0.00655 | 0.00519 | 0.00543 | 0.9573 | 0.72757 | 0.30493 | 3.37287 |
| Sedoheptulose-7-phosphate | 0.00623 | 0.0065 | 0.00852 | 0.00719 | 0.00621 | 0.00732 | 0.00673 | 0.00762 | 0.00665 | 0.00716 | 0.00693 | 0.0071 | 0.97645 | 0.73293 | 0.29874 | 8.40756 |
| β-Hydroxypyruvic acid | N/A | 0.0607 | 0.05689 | N/A | 0.0551 | 0.06823 | 0.06382 | 0.0552 | 0.0381 | 0.05116 | 0.05756 | 0.0553 | 1.04088 | 0.76141 | 0.29598 | 16.3712 |
| L-Homoserine/L-Threonine | 0.00133 | 0.00081 | 0.00152 | 0.00158 | 0.00138 | 0.001 | 0.0013 | 0.00127 | 0.0014 | 0.00139 | 0.00133 | 0.00127 | 1.0411 | 0.74184 | 0.28848 | 6.55927 |
| Glucosamine 6-phosphate | 0.04715 | 0.01104 | 0.00935 | 0.01724 | 0.01775 | 0.03105 | 0.018 | 0.02275 | 0.01911 | 0.02392 | 0.02051 | 0.02297 | 0.89286 | 0.74262 | 0.28758 | 20.6738 |
| PalCOA | 0.00024 | 0.0002 | 8.2E-05 | 0.00075 | 0.00052 | 0.0002 | 0.00028 | 0.00039 | 0.00078 | 0.0004 | 0.00036 | 0.00041 | 0.87805 | 0.75836 | 0.26952 | 9.19339 |
| Malic acid | 0.03611 | 0.01222 | 0.01648 | 0.02243 | 0.01934 | N/A | 0.03116 | 0.01676 | 0.02535 | 0.01851 | 0.02132 | 0.02295 | 0.929 | 0.77345 | 0.26746 | 12.0874 |
| α-muricholic acid | 0.05233 | 0.11428 | 0.08242 | 0.13837 | 0.14243 | 0.14587 | 0.11342 | 0.13119 | 0.08096 | 0.09107 | 0.10597 | 0.1125 | 0.94193 | 0.7634 | 0.26376 | 21.1558 |
| Trehalose 6-phosphate | 0.02946 | 0.00706 | 0.00575 | 0.00832 | 0.00973 | 0.01163 | 0.00921 | 0.01068 | 0.00946 | 0.01244 | 0.01206 | 0.01068 | 1.12916 | 0.76398 | 0.26311 | 13.3172 |
| Histidinol | 0.00667 | 0.00216 | 0.00249 | 0.00288 | 0.00457 | 0.0027 | 0.00415 | 0.00184 | 0.00496 | 0.00362 | 0.00375 | 0.00345 | 1.0867 | 0.77263 | 0.25323 | 24.1783 |
| Glyceric acid | 0.00427 | 0.00887 | 0.00756 | 0.00789 | 0.00804 | 0.00702 | 0.00804 | 0.00654 | 0.00834 | 0.00796 | 0.00732 | 0.00758 | 0.96608 | 0.77354 | 0.25221 | 21.9093 |
| 3-Phosphoglyceric acid | 0.04261 | 0.01409 | 0.00853 | 0.03082 | 0.02581 | 0.02632 | 0.01689 | 0.03109 | 0.02273 | 0.03482 | 0.02437 | 0.02637 | 0.92424 | 0.77698 | 0.24829 | 29.9892 |
| meso-Erythritol | 0.00328 | 0.00262 | 0.00148 | 0.00225 | 0.00243 | 0.00193 | 0.00166 | 0.00237 | 0.00282 | 0.00275 | 0.00241 | 0.00231 | 1.04601 | 0.78064 | 0.24413 | 9.18823 |
| 1-Oleoyl-rac-glycerol | 0.0008 | 0.00069 | 0.0009 | 0.00091 | 0.00087 | 0.00087 | 0.00118 | 0.00111 | 0.00042 | 0.00079 | 0.00083 | 0.00087 | 0.95577 | 0.79209 | 0.23114 | 21.668 |
| 12-Hydroxydodecanoic acid | 0.20508 | 0.18121 | 0.42597 | 0.1943 | 0.15983 | 0.21479 | 0.24074 | 0.22905 | 0.1689 | 0.24497 | 0.23328 | 0.21983 | 1.06185 | 0.7953 | 0.22751 | 7.54875 |
| γ-Linolenic acid | 0.05173 | 0.0282 | 0.04307 | 0.07286 | 0.03228 | 0.03969 | 0.06172 | 0.05036 | 0.05113 | 0.03703 | 0.04563 | 0.04798 | 0.95087 | 0.80236 | 0.21952 | 18.2178 |
| IsovalCOA | 1.4E-05 | 4.5E-05 | 7.2E-06 | 4.6E-05 | 4.3E-05 | 5.6E-05 | 4.7E-05 | 2.2E-06 | 6.7E-05 | 3.8E-06 | 3.1E-05 | 3.5E-05 | 0.88416 | 0.80385 | 0.21784 | 25.6457 |
| Uridine 5'-diphosphogalactose | 0.80466 | 0.07611 | 0.07301 | 0.16015 | 0.17039 | 0.07948 | 0.23254 | 0.21557 | 0.18431 | 0.39512 | 0.25686 | 0.2214 | 1.16015 | 0.8161 | 0.20403 | 25.7129 |
| Uridine 5′-diphospho-N-acetylgalactosamine | 2.21746 | 0.29698 | 0.38933 | 0.49295 | 0.54866 | 0.55432 | 0.72894 | 0.5391 | 0.80523 | 0.88037 | 0.78908 | 0.70159 | 1.12469 | 0.8171 | 0.20291 | 10.3728 |
| Melatonin | 2.6E-05 | 1.9E-05 | 1.4E-05 | 8.8E-06 | 4E-05 | 2.9E-05 | 9.8E-06 | 8.9E-06 | 2.5E-05 | 4.4E-05 | 2.2E-05 | 2.3E-05 | 0.91958 | 0.83094 | 0.18736 | 28.7226 |
| Octanoyl-carnitine (Carnitine_C8) | 0.05789 | 0.04099 | 0.08092 | 0.05778 | 0.04517 | 0.05591 | 0.06122 | 0.05184 | 0.05539 | 0.05048 | 0.05655 | 0.05497 | 1.02874 | 0.83198 | 0.1862 | 8.49874 |
| trans-1,2-Cyclohexanediol | 0.0102 | 0.01033 | 0.01302 | 0.01229 | 0.0109 | 0.01041 | 0.01215 | 0.01006 | 0.01075 | 0.01256 | 0.01135 | 0.01119 | 1.01438 | 0.8347 | 0.18314 | 19.7287 |
| Corticosterone | 0.002 | 0.00122 | 0.00155 | 0.0018 | 0.00089 | 0.00072 | 0.00282 | 0.00164 | 0.00103 | 0.00085 | 0.00149 | 0.00141 | 1.05772 | 0.85569 | 0.15966 | 20.5524 |
| Saccharic acid | 0.00313 | 0.00173 | 0.0017 | 0.00088 | 0.00076 | 0.00194 | 0.00193 | 0.00163 | 0.00083 | 0.00145 | 0.00164 | 0.00156 | 1.05484 | 0.86043 | 0.15437 | 21.9221 |
| N-PRO | 0.00048 | 0.0004 | 0.00023 | 0.00098 | 0.00093 | 0.00063 | 0.00061 | 0.00072 | 0.00027 | 0.00065 | 0.0006 | 0.00057 | 1.05141 | 0.86579 | 0.1484 | 11.5428 |

| | | | | | | | | | | | | | | | | |
|---|---|---|---|---|---|---|---|---|---|---|---|---|---|---|---|---|
| Cholesteryl oleate | N/A | 12.8183 | 11.8217 | 14.737 | 12.3119 | 12.9415 | 10.5018 | 13.6684 | 12.9016 | 13.9508 | 12.9222 | 12.7928 | 1.01012 | 0.88827 | 0.13086 | 22.0171 |
| 3-Methylphenylacetic acid | 0.01792 | 0.0081 | 0.01207 | 0.0121 | 0.01099 | 0.01059 | 0.01106 | 0.01164 | 0.01396 | 0.01264 | 0.01224 | 0.01198 | 1.02146 | 0.88408 | 0.12805 | 15.2287 |
| Nicotine | 0.00597 | 0.00346 | 0.00384 | 0.00517 | 0.00259 | 0.00472 | 0.00402 | 0.00369 | 0.0036 | 0.00453 | 0.00421 | 0.00411 | 1.02357 | 0.88462 | 0.12745 | 11.7223 |
| Mannose | 0.00885 | 0.00067 | 0.00153 | 0.00117 | 0.00264 | 0.00621 | 0.00232 | 0.00126 | 0.00274 | 0.00359 | 0.00297 | 0.00322 | 0.92238 | 0.88807 | 0.12363 | 25.6502 |
| Seleno-methionine | 0.34162 | 0.28001 | 0.59869 | 0.38803 | 0.29673 | 0.33358 | 0.41924 | 0.43485 | 0.35366 | 0.32138 | 0.38102 | 0.37254 | 1.02275 | 0.89461 | 0.11636 | 6.22843 |
| Mesoxalic acid | 0.02773 | 0.01626 | 0.01967 | 0.01969 | 0.0201 | 0.02057 | 0.02165 | 0.0241 | 0.01721 | 0.01843 | 0.02069 | 0.02039 | 1.01465 | 0.89752 | 0.11313 | 16.003 |
| MalCOA | N/A | 1.4E-05 | 5E-06 | 4E-05 | 4.1E-05 | 3E-05 | 3.1E-05 | 2E-05 | 3.6E-05 | 4.3E-07 | 2.5E-05 | 2.4E-05 | 1.05697 | 0.90401 | 0.11234 | 18.6355 |
| Lumichrome | 0.0859 | 0.06285 | 0.07633 | 0.08414 | 0.06238 | 0.08052 | 0.06697 | 0.07379 | 0.07296 | 0.08084 | 0.07432 | 0.07502 | 0.99072 | 0.90534 | 0.10447 | 9.77022 |
| Pentadecanoic acid | 0.02529 | 0.00705 | 0.01081 | 0.01006 | 0.01034 | 0.0087 | 0.0141 | 0.01076 | 0.01019 | 0.01768 | 0.01271 | 0.01229 | 1.03425 | 0.90967 | 0.09967 | 15.6027 |
| p-Coumaric acid | 0.01096 | 0.00737 | 0.00831 | 0.00899 | 0.00633 | 0.00789 | 0.00803 | 0.0086 | 0.00853 | 0.00934 | 0.00839 | 0.00848 | 0.98996 | 0.92018 | 0.08805 | 15.1499 |
| Adenosine 5'-diphosphoribose | 0.00116 | 0.00141 | 0.0014 | 0.00036 | 0.00053 | 0.00048 | 0.00142 | 0.0014 | 0.00101 | 0.00041 | 0.00097 | 0.00094 | 1.03091 | 0.92728 | 0.08019 | 8.7083 |
| Tryptophan | 0.21137 | 0.02207 | 0.02757 | 0.03319 | 0.03236 | 0.1244 | 0.06063 | 0.02961 | 0.06243 | 0.06692 | 0.06531 | 0.0688 | 0.94931 | 0.93212 | 0.07485 | 16.7524 |
| N-Acetyl-L-cysteine | N/A | 8.1E-06 | 2.5E-05 | 6E-05 | 2.4E-05 | 7.3E-05 | 6.6E-06 | 2.6E-05 | 2.7E-05 | 2E-05 | 2.9E-05 | 3.1E-05 | 0.95744 | 0.93737 | 0.07323 | 26.6592 |
| Adenosine 5'-monophosphate | 1.60904 | 1.08149 | 1.88695 | 1.50909 | 1.40702 | 1.15353 | 1.57844 | 1.83363 | 1.32709 | 1.54022 | 1.49872 | 1.48658 | 1.00816 | 0.94647 | 0.05901 | 3.50162 |
| 2'-Deoxyguanosine 5'-monophosphate | 4.2E-05 | 6.1E-06 | 4.3E-05 | 6.8E-06 | 2.6E-05 | 5.8E-06 | 2.3E-05 | 3.9E-05 | 5.3E-05 | 6.3E-06 | 2.5E-05 | 2.6E-05 | 0.96983 | 0.95151 | 0.05344 | 24.1281 |
| Phosphonoacetic acid | 0.03525 | 0.00545 | 0.0051 | 0.00854 | 0.00844 | 0.02077 | 0.00866 | 0.01083 | 0.01098 | 0.01344 | 0.01256 | 0.01294 | 0.9705 | 0.95158 | 0.05336 | 19.0325 |
| GluCOA | 2E-06 | N/A | 4.9E-06 | 8.7E-06 | 6.6E-06 | 6E-07 | 9.3E-06 | 7.3E-06 | 8.8E-06 | 2.2E-06 | 5.6E-06 | 5.6E-06 | 0.98263 | 0.96817 | 0.03719 | 21.6518 |
| Farnesyl pyrophosphate | 0.00241 | 0.00087 | 0.00131 | 0.0009 | 0.00056 | 0.00106 | 0.00104 | 0.00141 | 0.0015 | 0.00098 | 0.00121 | 0.0012 | 1.01155 | 0.96859 | 0.03461 | 12.2198 |
| Aspartic acid | N/A | 0.23467 | 0.30868 | 0.40818 | 0.36752 | 0.54083 | 0.49826 | 0.49169 | 0.41189 | 0.71346 | 0.32976 | 0.53123 | 0.62076 | 0.01822 | 0.03067 | 12.9712 |
| Pterine | 0.00422 | 0.00195 | 0.00169 | 0.00191 | 0.01213 | 0.0075 | 0.00245 | 0.00324 | 0.00382 | 0.00523 | 0.00438 | 0.00445 | 0.98468 | 0.97583 | 0.02663 | 28.1371 |
| Uridine | 0.01054 | 0.00092 | 0.0013 | 0.00111 | 0.00095 | 0.0043 | 0.0023 | 0.00259 | 0.00245 | 0.0029 | 0.00296 | 0.00291 | 1.01987 | 0.97687 | 0.02548 | 20.1678 |
| O-Phosphorylethanolamine | N/A | 0.10619 | 0.11037 | 0.20957 | 0.20113 | 0.18393 | 0.18569 | 0.25468 | 0.21623 | 0.25781 | 0.15682 | 0.21967 | 0.71387 | 0.07926 | 0.01616 | 13.678 |
| N-Acetyl-DL-methionine | 2.1E-05 | N/A | 3.3E-06 | 1.5E-06 | N/A | 9.5E-06 | 6.1E-06 | 1.7E-05 | N/A | 2.3E-06 | 8.8E-06 | 8.7E-06 | 1.00292 | 0.99702 | 0.00409 | 11.3723 |
| Isoleucine | 4.40407 | 1.22003 | 1.23215 | 1.91538 | 2.40207 | 1.80178 | 3.05371 | 1.72838 | 1.31388 | 3.27734 | 2.23474 | 2.23502 | 0.99988 | 0.9997 | 0.00034 | 5.22667 |
